# Comprehensive mapping of sensory and sympathetic innervation of the developing kidney

**DOI:** 10.1101/2023.11.15.567276

**Authors:** Pierre-Emmanuel Y. N’Guetta, Sarah R. McLarnon, Adrien Tassou, Matan Geron, Sepenta Shirvan, Rose Z. Hill, Grégory Scherrer, Lori L. O’Brien

## Abstract

The kidney functions as a finely tuned sensor to balance body fluid composition and filter out waste through complex coordinated mechanisms. This versatility requires tight neural control, with innervating efferent nerves playing a crucial role in regulating blood flow, glomerular filtration rate, water and sodium reabsorption, and renin release. In turn sensory afferents provide feedback to the central nervous system for the modulation of cardiovascular function. However, the cells targeted by sensory afferents and the physiological sensing mechanisms remain poorly characterized. Moreover, how the kidney is innervated during development to establish these functions remains elusive. Here, we utilized a combination of light-sheet and confocal microscopy to generate anatomical maps of kidney sensory and sympathetic nerves throughout development and resolve the establishment of functional crosstalk. Our analyses revealed that kidney innervation initiates at embryonic day (E)13.5 as the nerves associate with vascular smooth muscle cells and follow arterial differentiation. By E17.5 axonal projections associate with kidney structures such as glomeruli and tubules and the network continues to expand postnatally. These nerves are synapsin I-positive, highlighting ongoing axonogenesis and the potential for functional crosstalk. We show that sensory and sympathetic nerves innervate the kidney concomitantly and classify the sensory fibers as calcitonin gene related peptide (CGRP)^+^, substance P^+^, TRPV1^+^, and PIEZO2^+^, establishing the presence of PIEZO2 mechanosensory fibers in the kidney. Using retrograde tracing, we identified the primary dorsal root ganglia, T10-L2, from which PIEZO2^+^ sensory afferents project to the kidney. Taken together our findings elucidate the temporality of kidney innervation and resolve the identity of kidney sympathetic and sensory nerves.

## Introduction

Major functions of the kidney include the maintenance of body fluid and electrolyte homeostasis and modulation of blood pressure. These functions are maintained by a variety of intrinsic cellular mechanisms as well as extrinsic input from the nervous system (May et al., 2010; McMahon, 2016; Osborn et al., 2021). Peripheral neurons originating from ganglia innervate the kidney and are predominantly unmyelinated C-fibers (Kneupfer and Schramm, 1987; Booth et al., 2015; Osborn et al., 2021), although some myelinated renal afferent fibers have been reported in the kidney, yet their identity remains unclear (Simon and Schramm, 1983; Kneupfer and Schramm, 1987; Stella and Zanchetti, 1991). Kidney innervation consists of afferent sensory nerves that influence cardiovascular function through the relay of chemical and mechanical information from the kidney to the central nervous system (CNS), or efferent sympathetic nerves regulating renal function through the CNS (Osborn et al., 2021). Precise innervation and modulation of nerve activity maintains kidney physiology while aberrant nerve function is associated with several kidney diseases (DiBona & Kopp, 1997; Malpas, 2010; Sata et al., 2018). Therefore, modulating nerve activity is widely seen as a potential therapy to address kidney-related physiological disorders (Bhatt et al., 2022). Yet, to achieve such precise neuromodulation, an accurate mapping of the kidney nerve origins and a complete understanding of their molecular identity is necessary. Many studies to date have focused on sympathetic nerves and their regulation of mature kidney physiology. Therefore, we still lack detailed information about how both the sensory and sympathetic innervation of the kidney is established and how nerves pattern to reach their target cells and establish crosstalk.

Sympathetic nerves participate in the control of several kidney functions including vascular tone and glomerular filtration rate, sodium and water reabsorption, and renin release (Denton et al., 2004; DiBona, 2000; Fujisawa et al., 2011). They also modulate processes such as the release of cytokines to mediate inflammatory responses (Banek et al., 2016; Osborn et al., 2021; Veelken et al., 2008; Xiao et al., 2015). Overactivity of kidney sympathetic nerves contributes to the development and maintenance of hypertension (Osborn et al., 2021; Sata et al., 2018). In contrast to the numerous functions for sympathetic nerves in both normal physiology and pathophysiology, our understanding of kidney sensory nerve anatomy, classification, and their physiological roles remain poorly understood (Osborn et al., 2021). Kidney sensory nerves have been thus far been classified as transient receptor potential cation channel subfamily V member 1-positive (TRPV1^+^) due to their response to capsaicin and have also been shown to produce the nociceptor-associated neuropeptides calcitonin gene-related peptide (CGRP) and substance P (Osborn et al., 2021; Tyshynsky et al., 2023). Sensory neurons innervating the kidney project to the dorsal horn and synapse onto interneurons that project to CNS sites regulating cardiovascular function and in turn feedback to modulate kidney sympathetic nerve activity (Kopp, 2015). Recently, studies have established the importance of mechanosensitive PIEZO2 sensory nerves in the urogenital system to regulate bladder and sexual function (Lam et al., 2023; Marshall et al., 2020). However, whether PIEZO2 sensory nerves are also present in the kidney is unknown. Lastly, a recent study published by Cheng et. al. show parasympathetic cholinergic (ChAT^+^ and VAChT^+^) nerves innervate the renal artery and pelvis with cholinergic ganglion found within the kidney nerve plexus (Cheng et al., 2022). Whether these axons project to the kidney cortex and associate with cells or structures within kidney has not been elucidated to date. Initial studies of renal denervation in hypertensive patients led to a decrease in blood pressure although further intensive trials reported that certain patients experienced a rise in arterial pressure (Krum et al., 2013, 2014; Osborn & Foss, 2017; Schlaich et al., 2009; Symplicity HTN-2 Investigators et al., 2010; Townsend et al., 2017). To date, this method of controlling hypertension remains an attractive option with additional trials being conducted. However, such procedures non-selectively target both sensory and sympathetic nerves, therefore understanding the full extent of kidney nerve activity may inform efforts to optimize renal denervation strategies.

Ganglia located externally send axonal projections to the kidney. It has been reported that sympathetic innervation arises from the celiac, superior mesenteric, aorticorenal ganglia, and partially from the paravertebral chain depending on the model system being used (Barajas et al., 1992; Bell & McLachlan, 1982; Ferguson et al., 1986, 1988; Kuo et al., 1982). In rat, studies have shown that kidney sensory neurons project from dorsal root ganglion (DRG) at level T6 to L4, the majority of which derive from DRG at the level of T12–L3 (Donovan et al., 1983; Weiss & Chowdhury, 1998). However, the origin of sensory axons which innervate the mouse kidney is unclear. Moreover, the process of kidney innervation during development, in any model, has not been detailed thoroughly aside from a few recent studies reporting the close association of nerves with the developing kidney arterial tree (Honeycutt et al., 2023; Tarnick et al., 2023). In this study, we provide detailed spatiotemporal mapping of mouse kidney sensory and sympathetic neurons across embryonic and early postnatal development, as well as their persistence in the adult. We establish when kidney innervation begins, origins in the DRG, the spatial organization and growth of axons, their association with kidney cells and structures, as well as the identity of the nerves across these different stages. Altogether, we have generated a rich resource of knowledge surrounding kidney innervation that will inform studies of both kidney development and physiological regulation.

## Results

### Spatiotemporal mapping of kidney innervation

Due to the lack of knowledge on how kidneys are innervated during development, we set out to determine when neuronal axons first reach the kidney and how this network subsequently expands. We generated spatiotemporal maps of kidney innervation throughout development by performing wholemount immunostaining followed by 3D light-sheet imaging of either whole urogenital systems (E12.5, E13.5) or individual kidneys (E14.5, E16.5, E18.5, P0, P7). Kidneys and urogenital systems were immunostained with the pan-neuronal marker α-tubulin class III (TUBB3, TUJ1) to visualize axons, the arterial marker α-smooth muscle actin (SMA), and the endothelial marker CD31. Our previous studies have shown that the nerves and vasculature are tightly aligned in the developing kidney, therefore we wanted to further examine the association of these two networks at the various developmental timepoints (Honeycutt et al., 2023). Male and female kidneys were both examined, with no significant differences observed, therefore the data presented is representative of either sex. At E12.5, no TUBB3^+^ nerves or SMA^+^ vessels could be identified in the kidney, which was discerned by cytokeratin staining of the ureteric tree (**Fig. 1A; Movie S1**). SMA^+^ cells were primarily localized to the descending aorta with no SMA^+^ vascular mural cells yet found in the kidney, correlating with previous investigations of kidney vascularization **(Fig. 1A)** (Daniel et al., 2018). Despite the lack of axons in the kidney, we did observe the presence of ganglia and postganglionic nerve fibers tracking with the descending aorta (**Fig. 1A**). As organogenesis continues, TUBB3^+^ nerve fibers navigate towards the kidney and begin innervation at E13.5, closely associating with SMA^+^ mural cells which mark the maturing renal arteries (**Fig. 1B, arrowheads; Movies S2,S3**) (Daniel et al., 2018; Honeycutt et al., 2023; Luo et al., 2023; Tarnick et al., 2023). Despite the low SMA^+^ vascular mural cell coverage at this timepoint, an extensive CD31^+^ endothelial network is established in the kidney by E13.5 (**Fig. 1B**).

**Figure 1.**
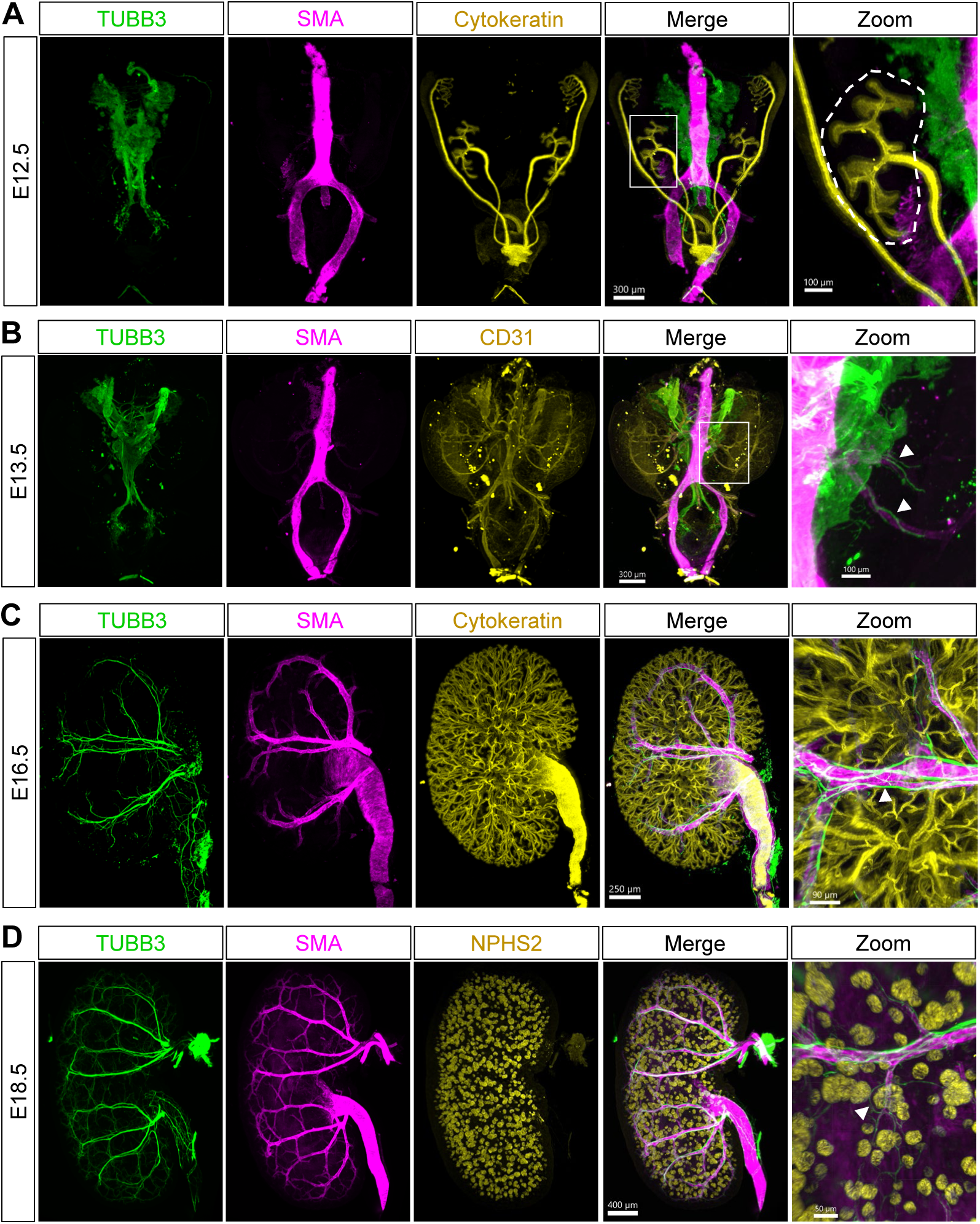
Kidney innervation commences at E13.5 as axons follow the differentiating arteries and continue to branch throughout development. **(A)** Pan-neuronal labeling (TUBB3, green) of an E12.5 urogenital system (UGS) shows that renal nerves are absent in the kidney at E12.5 as shown by immunostaining of the branching ureteric bud (cytokeratin, yellow). Ganglion and some postganglionic nerve fibers track along the descending aorta shown by smooth muscle actin (SMA, magenta). **(B)** Pan-neuronal labeling (TUBB3, green) of an E13.5 UGS shows that axons begin to innervate the kidney following the SMA+ vascular mural cells (magenta) of the developing arteries (white arrowhead, zoomed image). CD31 (yellow) highlights the extensive endothelial network that has formed but is yet to be coated with mural cells. **(C)** Axonal projections (TUBB3, green) continue to grow and branch following the developing arterial tree (SMA, magenta) at E16.5. Cytokeratin (yellow) marks the collecting duct system. Multiple axon bundles can be observed on an artery (white arrowhead, zoomed image). **(D)** By E18.5, axons (TUBB3, green) have undergone extensive growth with the arterial tree (SMA, magenta) and additional interstitial branching as the nerves navigate towards and around kidney structures such as glomeruli (NPHS2, yellow; white arrowhead, zoomed image) A minimum of n=3 kidneys were analyzed for each stage.

At E14.5, SMA coverage was more extensive, marking several arterial branches (**Fig. S1A**). Concurrently, the TUBB3^+^ nerve fibers continue tracking with the major arterial branches, only extending along the portion of endothelium with SMA^+^ vascular mural cell coverage (**Fig. S1A**). Additionally, axons track with the SMA^+^ ureteric smooth muscle layer (**Fig. S1A**). As development continues, axons grow and branch in coordination with the arterial network, highlighting a tight neurovascular interplay as the major arterial branches are established by E16.5 (**Fig. 1C; Movie S4**). Multiple nerve fibers associate with a single arterial branch and the larger fibers may consist of multiple axons which are tightly bundled (**Fig. 1C, arrowhead).** Once the major arterial branches were established, the axonal network expanded via interstitial branching as observed at E18.5 (**Fig. 1D; Movie S5**). The branching axons can be observed looping around structures such as glomeruli (**Fig. 1D, arrowhead; Movie S5**). Subsequent imaging of the postnatal kidney at postnatal day (P)0 (**Fig. S1B**) and P7 (**Fig. S1C**) revealed a significant increase in nerve fiber density and arborization of the network due to continued axonal growth and branching. Together, these data highlight the coordinated growth of an extensive neuronal network as kidney organogenesis progresses.

To visualize the innervation of target cells and structures within the developing kidney at greater resolution, we performed confocal imaging on E18.5 kidney sections. Supporting our wholemount 3D analyses, we observed the axons tightly associating with SMA+ vasculature, with several fibers often being associated with any one arterial branch (**Fig. 2A**). The nerves track with renal arteries and arterioles into the renal cortex and can separate from the SMA^+^ vascular mural cells to loop around and associate with the Bowman’s capsule of glomeruli (**Fig. S2B, Fig. 1D**) and tubule epithelia (**Fig. S2C**). Overall, kidney innervation is tightly linked to arterial differentiation with axons dissociating from the SMA^+^ vessels to navigate towards or around kidney target structures such as glomeruli.

**Figure 2:**
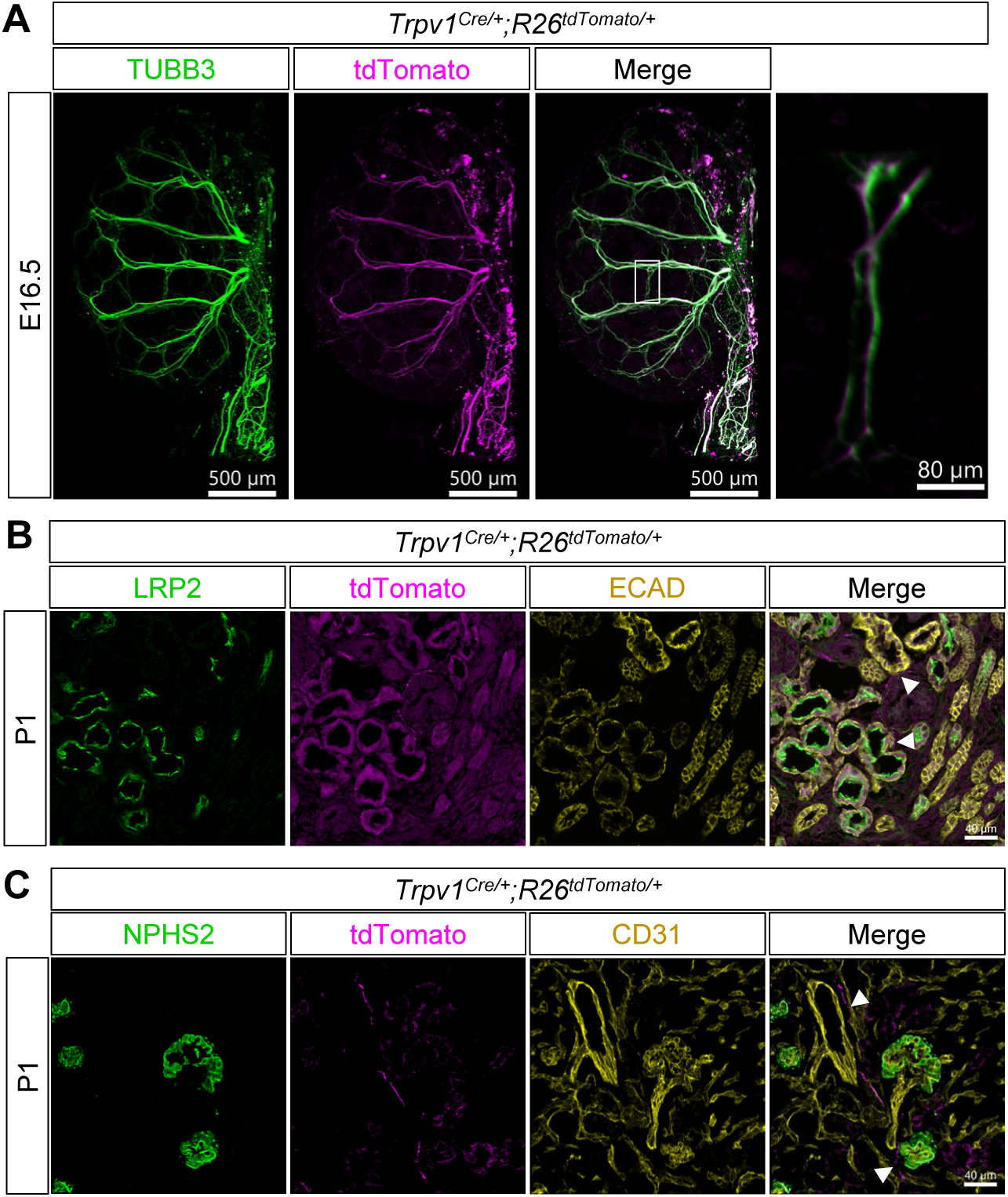
Nociceptive sensory afferent nerves innervate the developing kidney. **(A)** Wholemount immunostain of an E16.5 *Trpv1^Cre/+^;R26^tdTomato/+^* kidney showing tdTomato+ (magenta) axons colocalizing with the pan-neuronal marker TUBB3 (green). A magnified inset from the merged shows tdTomato^+^ axons overlapping with TUBB3. **(B)** *Trpv1^Cre/+^;R26^tdTomato/+^* P1 kidney section showing native tdTomato signal (magenta) with immunostained proximal tubules (LRP2, green) and epithelium (ECAD, yellow). TdTomato+ axons navigate around and closely associate with these structures (white arrowheads). **(C)** *Trpv1^Cre/+^;R26^tdTomato/+^* P1 kidney section showing native tdTomato signal (magenta) with immunostained podocytes (to mark glomeruli, NPHS2, green) and endothelium (CD31, yellow). TdTomato+ axons extend along the vasculature and associate with glomeruli (white arrowhead). A minimum of n=3 kidneys were analyzed for each stage.

### Sensory nerves innervate the developing kidney

While we have mapped the spatiotemporal innervation of the developing kidney, TUBB3 is a pan-neuronal marker labeling all nerve fiber types. It is well established that the mature kidney is innervated by sympathetic neurons that modulate renal physiology, however, the contribution of sensory nerves and the timing of sensory innervation in the kidney is poorly understood (Kopp, 2015; Osborn et al., 2021). To visualize the distribution and abundance of sensory neurons in the developing kidney, we crossed the *Trpv1^Cre^* line to the *Rosa(R)26^tdTomato^* reporter line (*Trpv1^Cre/+^;R26^tdTomato/+^*) to label all nociceptive TRPV1^+^ sensory neurons. TRPV1^+^ nociceptive neurons are thought to comprise the majority of, if not all sensory nerves within the kidney and therefore labeling this population should accurately represent the sensory contribution (Stocker & Sullivan, 2023; Tyshynsky et al., 2023). We performed wholemount imaging on *Trpv1^Cre/+^;R26^tdTomato/+^* kidneys at E16.5 to visualize the total complement of sensory nerve fibers traced from the start of kidney innervation. TdTomato^+^ signal extensively overlapped with the pan-neuronal marker TUBB3 (**Fig. 2A**). To visualize these sensory fibers and their association with kidney structures in greater detail, we examined sections from *Trpv1^Cre/+^;R26^tdTomato/+^* kidneys at P1. The tdTomato^+^ sensory nerves are found adjacent to LRP2^+^ proximal tubules and E-cadherin (ECAD)^+^ epithelial structures (**Fig. 2B**). Additionally, sensory axons are found adjacent to arteries, arterioles, and glomeruli (**Fig. 2C**). Taken together, these findings show that sensory nerves make up a significant portion of the neuronal network in the developing kidney and associate with kidney structures by early postnatal stages.

Nociceptive sensory afferent nerve identity and function can also be defined by the production of neuropeptides such as calcitonin gene-related peptide (CGRP) and substance P. To assess renal peptidergic sensory innervation, we immunostained kidney sections at E14.5, E15.5, E18.5, P0, and adult stages using antibodies for CGRP and substance P. At E14.5 we could not yet detect CGRP^+^ or substance P^+^ nerves (**Fig. S3A, S3B**). However, by E15.5 we observed CGRP^+^ sensory nerves that closely associated with renal arteries (**Fig. 3A**). To circumvent cross-reactivity issues due to several of our antibodies being raised in the same species, we crossed the transgenic podocin Cre line (*Pod-Cre^tg^*) to the *R26^tdTomato^* reporter (*Pod-Cre^tg/+^;R26^tdTomato/tdTomato^*) thereby labeling all glomerular podocytes with tdTomato. Assessing kidneys from these animals at P0, we found that CGRP^+^ sensory nerves came in close proximity to glomeruli and proximal tubules (**Fig. 3B**). CGRP^+^ sensory neurons were also found in adult kidneys, where they similarly associated with the vasculature and Bowman’s capsule of glomeruli (**Fig. 3C**). In contrast to CGRP, substance P immunolabelling was not detected until E18.5 where these sensory nerves were observed adjacent to blood vessels and glomeruli (**Fig. 3D**). In the adult kidneys, substance P maintained this localization pattern and these fibers were similarly found associated with the vasculature and Bowman’s capsule (**Fig. 3E**). Peripheral neurons innervating the kidney are predominantly unmyelinated C-fibers (Kneupfer and Schramm, 1987; Booth et al., 2015; Osborn et al., 2021). Yet, it has been suggested that some myelinated afferent fibers are present in the kidney (Simon and Schramm, 1983; Kneupfer and Schramm, 1987; Stella and Zanchetti, 1991), although it was unclear if these fibers were peptidergic. To address this, we assessed the colocalization of CGRP+ fibers and Neurofilament heavy polypeptide (NFH), a component of large diameter myelinated axons, in adult kidneys (**Fig. S4**). In the kidney cortex, a few NFH+ fibers were observed although they did not come in proximity to glomeruli and CGRP signal did not overlap with the fibers. In the renal pelvis, we observed minor overlap of NFH^+^ and CGRP+ along the pelvic urothelium **(Fig. S4)**, supporting the presence, although sparse, of myelinated sensory fibers within the kidney.

**Figure 3:**
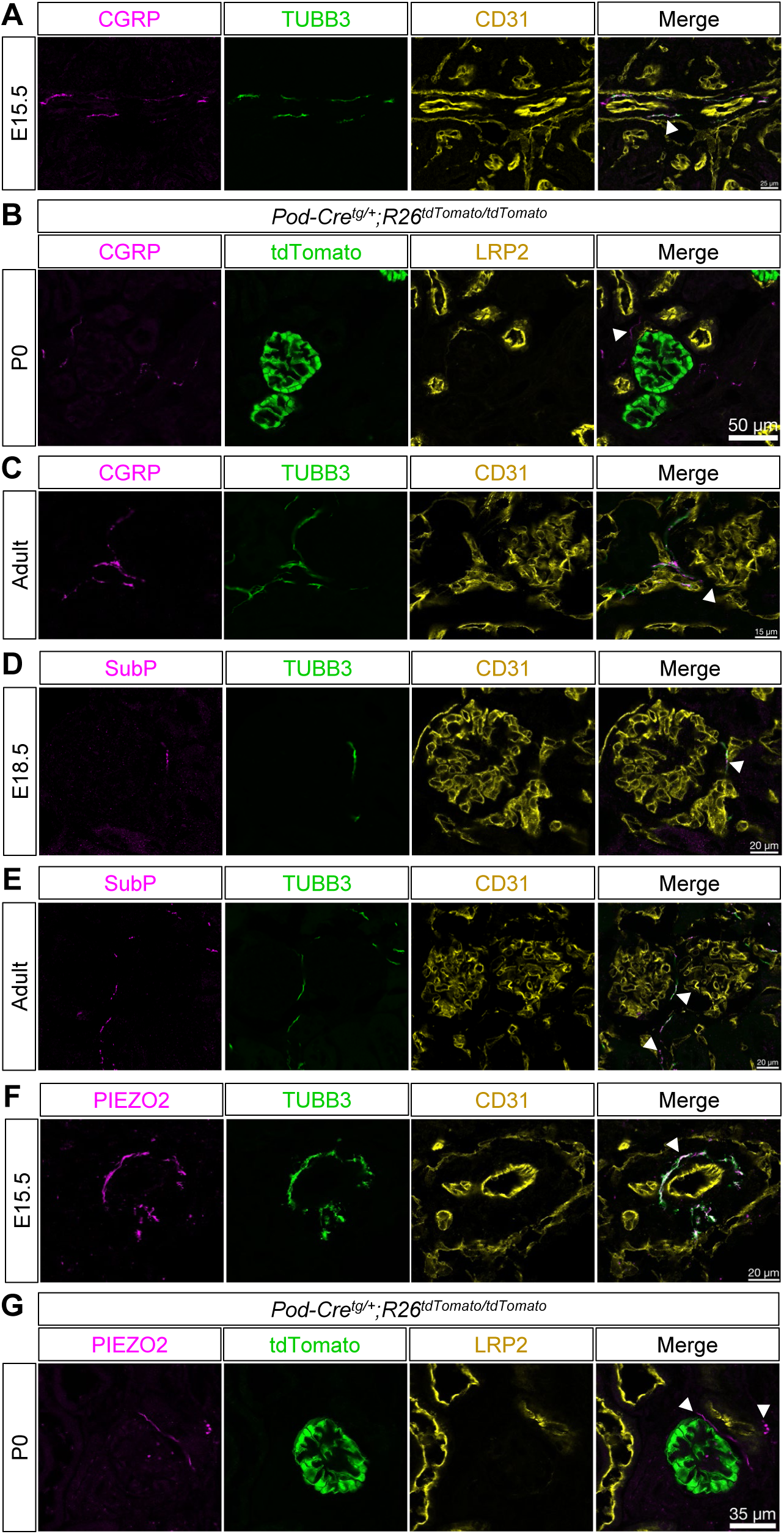
Peptidergic and mechanosensory afferent nerves are present in the developing kidney. **(A)** E15.5 kidney section immunostained for the neuropeptide CGRP^+^ (magenta) and pan-neuronal marker TUBB3 (green). CGRP^+^ nerves mainly associate with the kidney vasculature (CD31, yellow; white arrowhead) at this stage. **(B)** Sections of *Pod-Cre^tg/+^;R26^tdTomato/tdTomato^* P0 kidneys immunostained for CGRP^+^ (magenta). CGRP+ axons are found near (white arrowhead) glomeruli (tdTomato, green) and close to proximal tubules (LRP2, yellow). **(C)** Adult kidney section immunostained for CGRP (magenta) and colocalized with TUBB3 (green) shows CGRP^+^ axons closely associating with the vasculature (CD31, yellow) and in close proximity to glomeruli (white arrowhead). **(D)** E18.5 kidney section immunostained for substance P (SubP, magenta) shows overlap with TUBB3 (green). SubP^+^ nerves were found associated with the vasculature (CD31, yellow, white arrowhead). **(E)** Adult kidney section immunostained for SubP^+^ (magenta) and TUBB3 (green) shows SubP+ nerves along the vasculature (CD31, yellow) and in close proximity to glomeruli (white arrowhead). **(F)** E15.5 kidney section immunostained for PIEZO2 (magenta) and TUBB3 (green) show PIEZO2+ nerves associating with the vasculature (CD31, yellow; white arrowhead). **(G)** Sections of *Pod-Cre^tg/+^;R26^tdTomato/tdTomato^* P0 kidneys immunostained for PIEZO2 (magenta) show PIEZO2+ nerves closely associating (white arrowheads) with glomeruli (tdTomato, green) and proximal tubules (LRP2, yellow). A minimum of n=3 kidneys were analyzed for each stage.

PIEZO2 is a mechanoreceptor utilized by a subset of peripheral sensory neurons (Coste et al., 2010). PIEZO2^+^ neurons innervate organs of the urogenital system such as the bladder to help coordinate urination and genital sub-regions to support sexual function (Lam et al., 2023; Marshall et al., 2020). Therefore, we used immunofluorescence staining of PIEZO2 on kidney sections to examine whether PIEZO2^+^ neurons also innervate the kidney during development. At E15.5, we observed PIEZO2^+^ neurons associating with CD31^+^ large vessels (**Fig. 3F**). By P0, we found PIEZO2^+^ neurons adjacent to glomeruli and proximal tubules (**Fig. 3G**). Collectively, our findings demonstrate the diversity of kidney sensory innervation which consists of both nociceptive and mechanosensitive neurons. The timing of when these neurons and associated neuropeptides can be detected varies throughout kidney development. However, all can be detected by birth and follow a similar pattern of association with the renal vasculature, glomeruli, and tubules.

### Retrograde tracing identifies the DRG origin of kidney sensory neurons

The somata of peripheral sensory neurons are located in DRG present along the spinal column. To identify the DRG that contain the sensory neurons projecting to the kidney, we performed retrograde tracing experiments. We injected into the right kidney (ipsilateral) of adult mice the fluorescently labeled neuronal tracer Cholera toxin subunit B (CTB). Four days after the injections, both the injected and non-injected kidney as well as the ipsilateral and contralateral DRG were dissected to determine the origin of kidney sensory innervation (**Fig. 4A**). By examining the fluorescent signal from all cervical, thoracic, lumbar, and sacral DRGs, we determined that kidney sensory innervation traces to the ipsilateral DRG of the lower thoracic (T10-T13), lumbar (L1-L6), and sacral (S1-S2) regions (**Fig. 4B**). However, the majority of sensory innervation projected from the lower thoracic (T10-T13) and upper lumber (L1-L2) regions (**Fig. 4B**). Ipsilateral kidney sections showed CTB^+^ nerves near vasculature and glomeruli (**Fig. 4C**). Strikingly, we observed CTB^+^ cells within the contralateral DRG of the upper lumbar region (L1-L2), suggesting these DRG may also innervate the kidney (**Fig. 4B**). Additionally, we observed a few CTB^+^ nerve fibers around blood vessels in the contralateral kidney (**Fig. 4C**). No difference was identified between male and female mice in the tracing results obtained.

**Figure 4:**
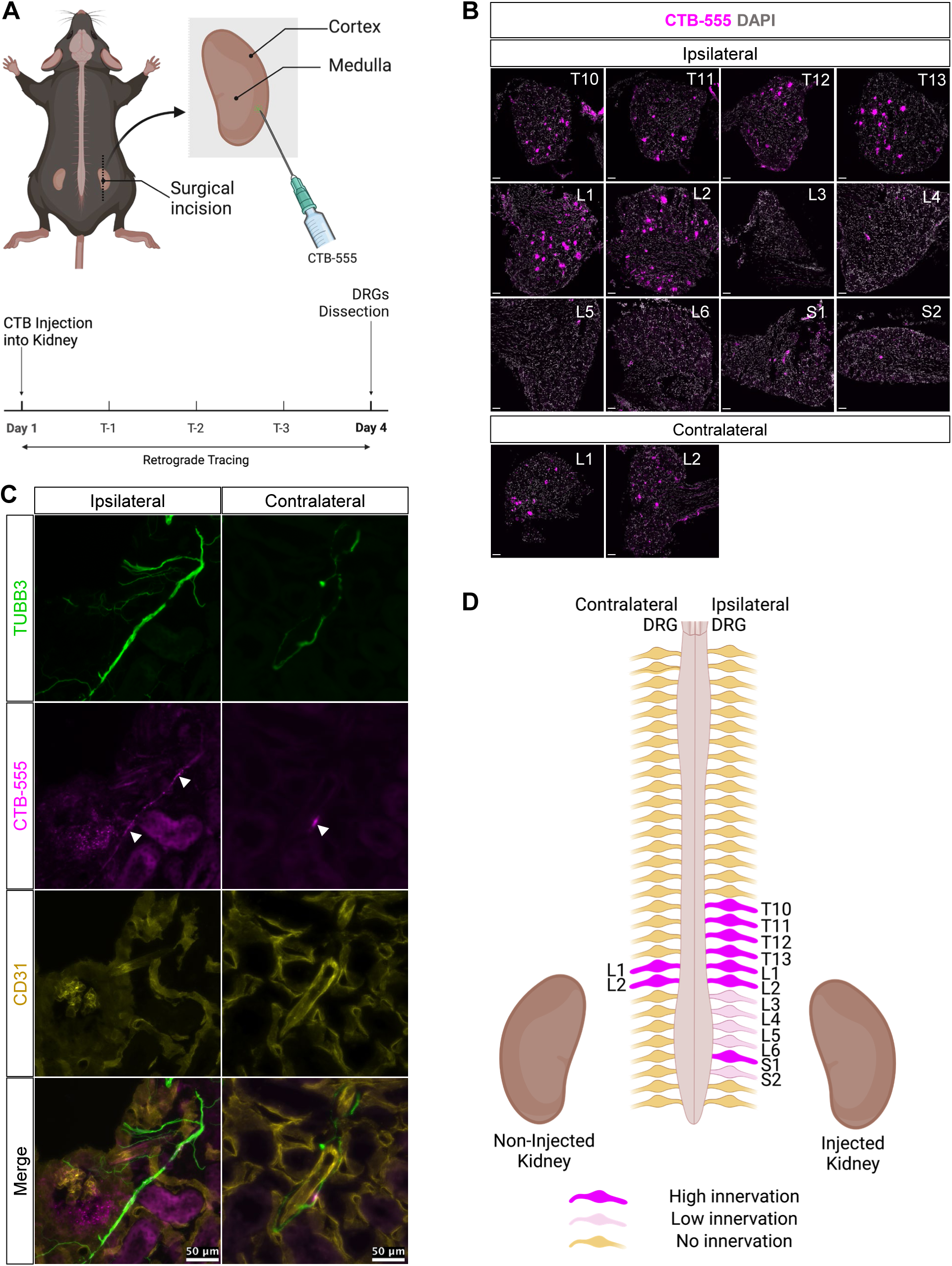
CTB retrograde tracing shows axon trajectories and labeling of thoracic, lumbar, and sacral DRGs. **(A)** Schematic of retrograde tracing injection and timeline. CTB-555 was injected into the right kidney of adult mice. Three days following the injection both kidneys were dissected as well as the ipsilateral DRGs (injected side) and contralateral DRGs (non-injected side). **(B)** Sensory innervation primarily traces in the ipsilateral DRGs to the lower thoracic region (T10-T13), lumbar region (L1-L6), and sacral region (S1-S2). Tracing to the contralateral DRGs was observed in the upper lumbar region of L1 and L2. n=4 males and n=4 females, Scale bar =50μm. **(C)** Ipsilateral and contralateral kidney sections of CTB-injected kidneys. Immunostaining of the ipsilateral kidney section showed CTB^+^ staining (magenta) colocalizing with TUBB3 (green) near glomeruli (CD31, yellow, arrowhead) and vasculature (CD31, yellow). CTB^+^ staining (magenta, arrowhead) colocalized with TUBB3 (green) near vasculature (CD31, yellow) in the contralateral (non-injected) kidney. **(D)** Schematic representation of the spinal level and DRG origin of kidney sensory afferent innervation.

Next, we specifically traced PIEZO2^+^ sensory neurons using the recombinant adeno-associated virus *pAAV-Flex-tdTomato* that we injected into the right kidney of *Piezo2^EGFP-IRES-Cre/+^* mice (**Fig. S5A**). The tracing confirmed innervation of the kidney by PIEZO2^+^ sensory neurons and shows that the majority of PIEZO2^+^ nerves project from the lower thoracic (T10-T13) and lumbar (L1 and L2) DRG (**Fig. S5B,C**). Taken together, the retrograde tracing experiments validated sensory innervation of the kidney and uncovered their origins within the lower thoracic, lumbar, and sacral regions (**Fig. 4D, S5C**). The observation that CTB^+^ cells were found in L1 and L2 contralateral DRG, along with the presence of labeled axons in the contralateral kidney, suggests the possibility of cross communication between the kidneys and DRG on the opposite sides of the body.

### Sympathetic and cholinergic innervation of the developing kidney

Sympathetic postganglionic neurons produce tyrosine hydroxylase (TH), an enzyme involved in catecholamine synthesis, and this marker has previously been utilized in the adult kidney to localize sympathetic nerves (Tiniakos et al., 2004; Torres et al., 2021). Thus, to determine the distribution of renal sympathetic nerves in the developing kidney we performed wholemount immunolabeling with a TH antibody at E14.5 and P0 (**Fig. 5A,B**). At E14.5, TH^+^ nerve fibers were detected in the kidney, overlapping with TUBB3 and tracking with the arteries (**Fig. 5A**). By P0 the sympathetic nerves had branched significantly with the arterial tree and the axons were found navigating around structures such as LRP2^+^ proximal tubules (**Fig. 5B**). We also examined kidney sections and similarly observed TH^+^ axons primarily associated with renal arteries at earlier stages of development (**Fig. 5C**). By P0, the nerves could be found abutting glomeruli and proximal tubules (**Fig. 5D**). We further validated sympathetic nerve identity by co-immunostaining adult kidney sections with the sympathetic marker neuropeptide Y (NPY) and TH. All NPY+ fibers examined, such as those adjacent to glomeruli, were TH+ confirming their sympathetic identity (**Fig. 5E**). We also investigated the overlap of TH+ and CGRP+ fibers and found them co-associating with glomeruli and in the renal pelvis **(Fig. S6A)**. We then quantified the percent of glomeruli that were innervated by either TH+ fibers, CGRP+ fibers, or both. A previous study performed similar quantitation and found CGRP+ fibers associated with ∼48% of glomeruli in wild type adult mice (Tyshynsky et al., 2023). Corroborating these findings, our analyses identified 40% of glomeruli were associated with CGRP+ fibers. TH+ axons associated with 50% of glomeruli and ∼37% were innervated by both CGRP+ and TH+ axons. **(Fig. S6B)**. Taken together, these findings show that sympathetic neurons innervate the kidney early in development, tracking with the vasculature and colocalizing with CGRP+ fibers as they make their way to nephron targets such as tubules and glomeruli in alignment with their known physiological functions (DiBona, 2004; DiBona & Kopp, 1997; Osborn & Foss, 2017; Sata et al., 2018).

**Figure 5:**
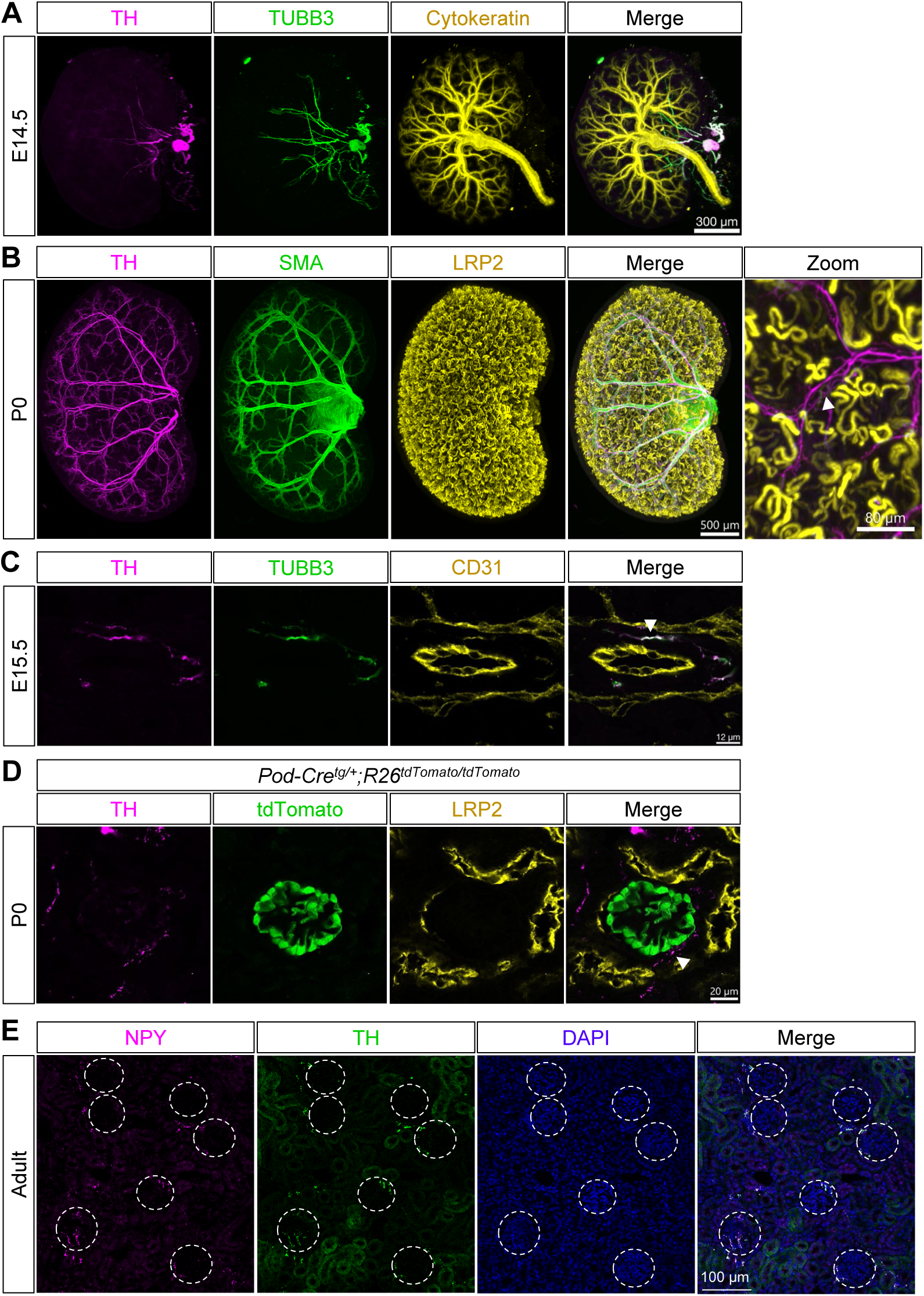
Noradrenergic sympathetic nerves innervate the developing kidney. **(A)** E14.5 wholemount immunostained kidney labeled with tyrosine hydroxylase (TH, magenta) shows that the first innervating axons (TUBB3, green) are TH+. Cytokeratin (yellow) marks the developing collecting duct system. **(B)** P0 wholemount immunostained kidney labeled with TH (magenta) shows that the TH+ nerves have branched extensively alongside the arterial tree (SMA, green). LRP2 marks the proximal tubules (yellow) and the magnified inset shows TH^+^ nerves (magenta) navigating around and in close proximity to proximal tubules (white arrowheads). **(C)** E15.5 kidney section immunostained for TH (magenta) and TUBB3 (green) shows TH+ axons around the vasculature (CD31, yellow; white arrowhead). **(D)** Sections of *Pod-Cre^tg/+^;R26^tdTomato/tdTomato^* P0 kidneys immunostained for TH (magenta) show TH^+^ axons near glomeruli (tdTomato, green; white arrowhead) and proximal tubules (LRP2, yellow). A minimum of n=3 kidneys were analyzed for each stage. **(E)** Image of the adult kidney cortex immunostained for the neuropeptide NPY (magenta), TH (green), and the nuclear counterstain DAPI (blue). Glomeruli are encircled in white dotted lines. No TH^+^/NPY^-^ fibers were observed. Image is representative of n=2 male mice.

Previous studies showed that during development, TH may also be produced by cholinergic parasympathetic nerves (Howard, 2005). A recent study demonstrated parasympathetic innervation of the main renal artery and pelvis, yet there remains no anatomic evidence of significant parasympathetic innervation into the kidney. To identify whether there are cholinergic nerves in the developing kidney, we immunostained sections at E14.5, E15.5, P0, and adult for the cholinergic marker vesicular acetylcholine transporter (VAChT) (**Fig. S7**). We did not observe VAChT^+^ nerves overlapping with pan-neuronal marker TUBB3 at E14.5 (**Fig. S7A**) even though the first nerve fibers innervating the kidney at E14.5 were TH^+^ (**Fig. 5A**). By E15.5, we observed that TUBB3^+^ nerves overlapped with VAChT signal, and these nerves were primarily associated with the renal artery in the kidney shown by CD31 immunostaining (**Fig. S7B**). Vascular-associated innervation increased by E16.5, and we observed VAChT^+^ fibers around proximal tubules and glomeruli at P0 (**Fig. S7C**). In adults, the VAChT^+^ fibers were found associated with the kidney vasculature (**Fig. S7D**). In sum, these data lend support to the presence of parasympathetic innervation into the kidney with axons reaching nephrons.

### Evidence for the establishment of neuroeffector junctions during kidney development

In order to exert an impact on kidney development and physiology, neurons need to establish functional crosstalk with kidney cells. To investigate the formation of neuroeffector junctions, synapse-like structures that form between neurons and non-neuronal cells, we characterized the cellular localization of a synaptic marker by immunofluorescence on cryosectioned kidneys. Using an antibody raised against synapsin I (SYN1), a membrane-associated phosphoprotein contained within small synaptic vesicles of presynaptic nerves, we investigated if and at which stage SYN1 is detected. At E14.5 SYN1^+^ axons were observed around the vasculature (**Fig. 6A**). At E16.5, there was similar localization of SYN1^+^ nerves mainly associated with the vasculature (**Fig. 6A**). By P0, SYN1^+^ axons were localized near proximal tubules and glomeruli (**Fig. 6B**). A similar pattern was retained in the adult with SYN1^+^ axons found along the vasculature and adjacent to glomeruli (**Fig. 6C**).

**Figure 6:**
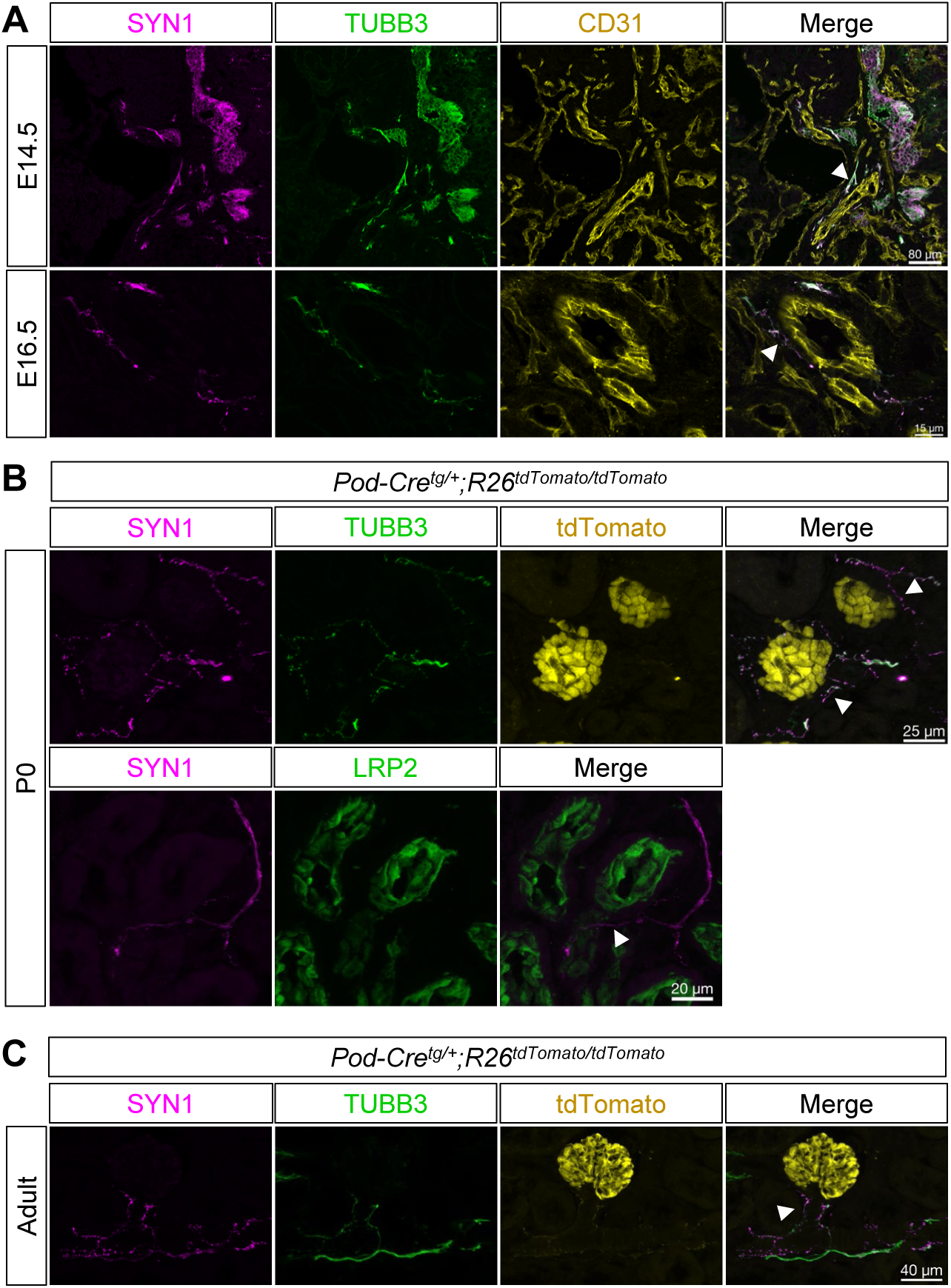
Nerves form putative neuroeffector-junctions with kidney targets during development. **(A)** Immunostaining of E14.5 kidney sections for synapsin I (SYN1, magenta) shows overlap with TUBB3 (green) and association of axons with the main artery (CD31, yellow; white arrowhead). **(B)** E15.5 kidney sections immunostained for SYN1 (magenta) and TUBB3 (green) similarly show SYN1^+^ nerves positive around the vasculature (CD31, yellow). **(C)** Sections of *Pod-Cre^tg/+^;R26^tdTomato/tdTomato^* P0 kidneys immunostained for SYN1 (magenta) show SYN1+ axons (TUBB3, green) near glomeruli (tdTomato, green; white arrowheads, top panel) and proximal tubules (LRP2, yellow; White arrowhead, bottom panel). **(D)** Sections of *Pod-Cre^tg/+^;R26^tdTomato/tdTomato^* adult kidneys immunostained for SYN1 (magenta) show SYN1+ axons (TUBB3, green) along the arterioles near glomeruli (tdTomato, green; white arrowhead). A minimum of n=3 kidneys were analyzed for each stage.

Next, we investigated the developmental expression of receptors associated with various neuronal signaling mechanisms and known post-synaptic markers to understand which kidney cells are responsive to neuronal signaling. We utilized a publicly available single cell RNA-seq dataset from the E18.5 kidney to obtain the enrichment of these factors in the various cells of the developing kidney (Combes et al., 2019) (**Fig. S8**). These data show that a variety of cell types and structures including nephron progenitors, stromal cells, renal vesicles, and the S-shaped body express α-adrenergic receptors *Adra1a* and *Adra1b*, whereas *Adra2a* and *Adra2b* were enriched in the distal tubules, ureteric epithelium, and proximal tubules (**Fig. S8A**). We also noted the β-adrenergic receptor *Adrb2* was strongly enriched in the resident immune cells while *Adrb3* was expressed by stromal cells (**Fig. S8A**). Next, we also looked for the expression of major purinergic receptors in the developing kidney. Many of the ATP and Adenosine receptors including *P2rx4, P2rx7, P2ry6, P2yr12, P2yr13, P2yr14, Lpar6,* and *Lpar4* showed enrichment in the resident immune cells. Endothelial cells, tubule epithelium, and ureteric epithelium showed expression of *Ador2a, P2ry1, Lpar6, Lpar4,* and *Adora1* while *P2rx4* and *P2ry14* were moderately enriched in podocytes (**Fig. S8B**). Lastly, we observed the enrichment of known post-synaptic markers or synapse-promoting markers like *Vamp2, Tenm3, Tenm4, Ephb1, Ephb3, Epha2, and Grm7* by podocytes, *Nlgn1, Epha2, Ephb1, and Ephb4* by endothelial cells, and *Gria3, Ephb2, Ephb3, Ephb4, Ephb6, Epha1, Epha4, Epha7, Tenm2, Dlg4, Dlg1, Nlgn3* by the distal and proximal tubules (**Fig. S8C**). Collectively, the single-cell expression data provides evidence that cells within the developing kidney express a variety of receptors and post-synaptic markers suggesting they are competent to receive and respond to neuronal-derived signals.

### The spatial organization of nerves is conserved in the adult human kidney

Given the specific distribution of kidney nerves around important structures such as glomeruli and tubules in the mouse, we aimed to validate that innervation of structures in the human adult kidney was similar. First, we immunostained adult human cortex cryosections with sympathetic nerve marker TH. We observed TH^+^ axons looping around glomeruli and associating with the Bowman’s capsule similar to the mouse. These sympathetic nerves were also found near epithelial structures (ECAD+) (**Fig. 7A; Movie S6**). Next, we sought to confirm the presence of PIEZO2^+^ mechanosensory afferents in the adult human kidney. We identified PIEZO2^+^ nerves that were closely associated with glomeruli and vessels as well as the kidney epithelium (**Fig. 7B**). To validate that the nerves were actively signaling we assayed for the presence of neuroeffector junctions by immunostaining for SYN1. We observed SYN1^+^ nerve fibers looping around glomeruli and around tubules (**Fig. 7C, Movie S7**). Taken together, these data show that the close association of both sympathetic and sensory nerves with vessels and nephrons is conserved in humans.

**Figure 7:**
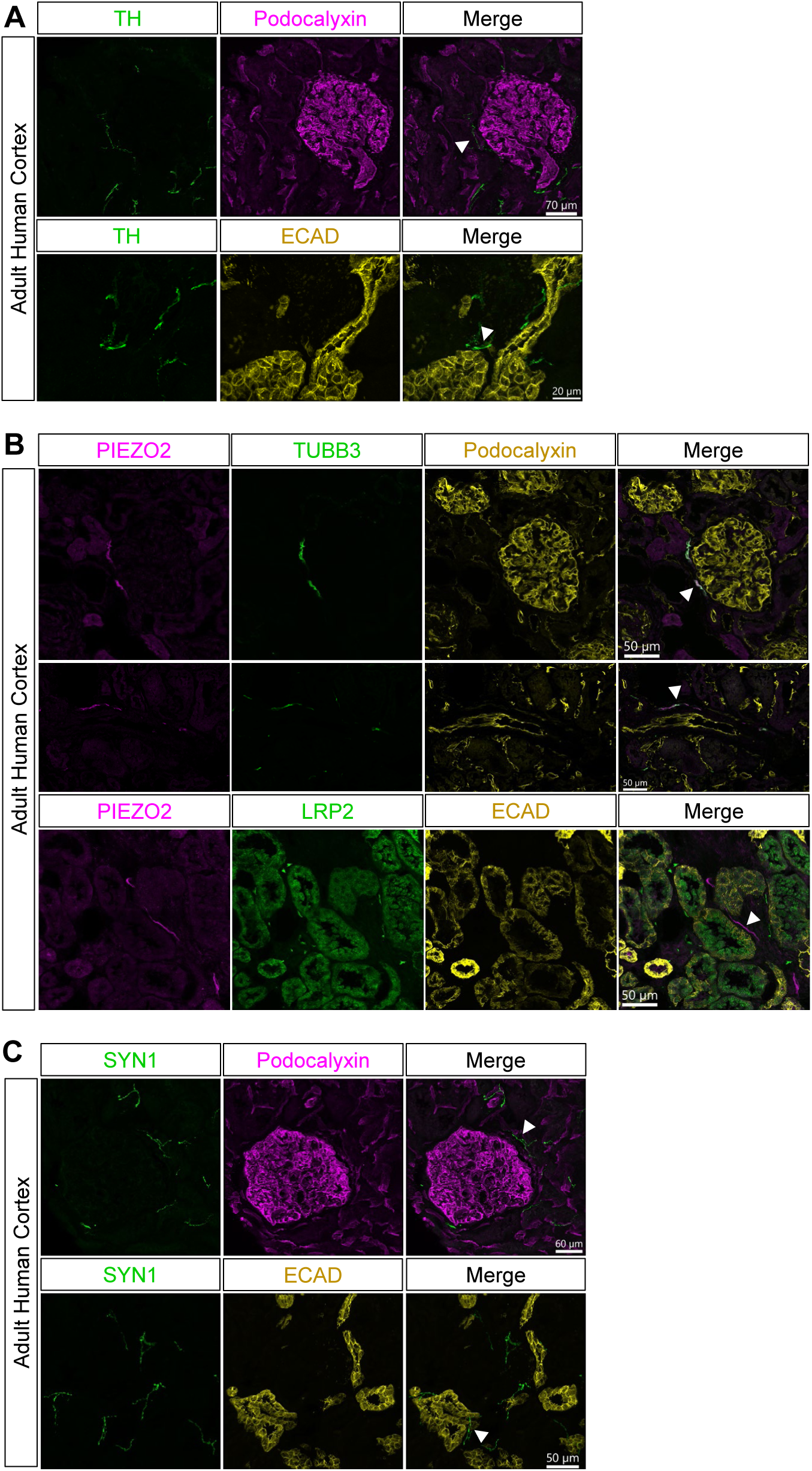
Sympathetic and sensory nerves show similar structural associations in the adult human kidney. **(A)** Adult human kidney sections immunostained for TH^+^ sympathetic nerves (green) can be observed associating with glomeruli shown by Podocalyxin (magenta; top panel, white arrowhead) and in close proximity to epithelial structures (ECAD, yellow; bottom panel, white arrowhead). **(B)** Adult human kidney sections immunostained for PIEZO2 (magenta) show PIEZO2^+^ overlapping with TUBB3 (green) associating with glomeruli (Podocalyxin, yellow; white arrowhead, top panel) and the vasculature (Podocalyxin, yellow; white arrowhead, middle panel). PIEZO2+ sensory nerves were also observed near epithelial structures (LRP, green; ECAD, yellow; white arrowhead, bottom panel). **(C)** Adult human kidney sections immunostained for SYN1 (green) show SYN1+ axons near glomeruli (Podocalyxin, magenta; white arrowhead, top panel) and epithelial structures (ECAD, yellow; white arrowhead, bottom panel).

## Discussion

In this study, we set out to investigate the process of kidney innervation across embryonic and postnatal mouse development. We generated spatiotemporal maps of innervation using markers specific to various classes of axons and identified the presence of both sensory and sympathetic nerves, as well as evidence for parasympathetic innervation. High resolution imaging of kidney sections provided insights into the association of nerves with kidney structures such as the vasculature, glomeruli, and tubules and the possible establishment of neuroeffector junctions. Finally, we validated that the human kidney shows similar associations of sympathetic and sensory nerve subclasses with nephron structures. Altogether, we have garnered substantial new insights into the process of kidney innervation with implications for both development and physiological regulation.

### The close association of nerves with kidney vasculature

We established that kidney innervation begins at E13.5 and coincides with the initiation of arterial differentiation as vascular smooth muscle cells associate with the nascent endothelium. In the developing kidney, it has been shown that arterial differentiation is first established at E13.5 and progresses toward the kidney periphery (Daniel et al., 2018; Luo et al., 2023). Our findings of a strict association of axons with vascular smooth muscle cells correlates with a recent description of kidney innervation as well as our own recent studies (Honeycutt et al., 2023; Tarnick et al., 2023). We hypothesize that SMA+ vascular mural cells may be signaling to the axons, thereby guiding the nerves as the arteries differentiate and grow. This has been shown in other systems with axons responding to vascular-derived cues like artemin (*Artn1*) and neurotrophin 3 (*Ntf3*) (Glebova & Ginty, 2005). In the kidney, *Ntf3* is expressed by nephron progenitor cells and early developing nephrons whereas *Artn1* does not appear to be expressed, as inferred from single cell RNA-seq data of the developing (E18.5) kidney (Combes et al., 2019). Whether *Ntf3* or *Artn1* are expressed by the vasculature at earlier stages and mediate kidney innervation not been investigated to date. Once established, continued axonal growth and survival relies on neurotrophic factors such as nerve growth factor (NGF). During nephrogenesis, NGF is produced by cortical, medullary, and ureteric stroma (Combes et al., 2019; Glebova & Ginty, 2004). Additionally, it was established that NGF is required for the complete sympathetic innervation of the kidney via its interaction with its innate receptor tropomyosin receptor kinase A (TRKA) (Glebova & Ginty, 2004). Moreover, proper innervation also requires chemotactic guidance cues released by adjacent cells within the tissue. In the kidney, we and others have established that proper patterning of the neurovascular network is mediated by the guidance cue netrin-1 (Honeycutt et al., 2023; Luo et al., 2023). However, we established that the endothelium is the primary responder to netrin-1 guidance cues, with the nerves following the vascular pattern, suggesting the SMA^+^ vascular mural cells themselves play a significant role in guiding kidney innervation (Honeycutt et al., 2023). While it has been established that netrin-1 plays a key role in synapse formation and axonal regeneration in the brain, we do not know whether the same is true in the kidney (Zheng et al., 2018). Additional studies are necessary to establish the full complement of cues necessary for proper kidney innervation and the establishment of functional crosstalk.

### The diversity of kidney nerves

During kidney development, we found that both sensory and sympathetic nerves innervate the tissue early in development and branch following the arterial tree. Sensory nerves could be classified as nociceptive by TRPV1^+^ lineage tracing. The limitation of this approach is that as neurons mature, they may not remain TRPV1^+^. As such, and with the lack of our ability to identify a TRPV1 antibody that worked on our kidneys, this lineage trace only serves as a proxy for TRPV1^+^ nerves that innervate the kidney from the onset. However, we were able to further classify the sensory neurons by their production of the neuropeptides CGRP and substance P. CGRP^+^ nerves were identified earlier in development than substance P^+^ nerves (E15.5 vs. E18.5), but both did not appear until innervation had already been established for several days. With the importance of neurotrophic factors for proper sensory afferent nerve function, these delays in production could be due to changes in molecular cues required for their specific growth and function (Ernsberger et al., 2020; Golden et al., 2010). The expression of their innate target receptor on kidney cells could also play an important role in triggering neuropeptide production. Despite these differences, all the sensory nerve markers showed similar tracking with the vasculature and association with kidney target structures such as glomeruli and tubules around E18.5 to P0.

Recent studies have established the expression of the *Piezo2* mechanosensory receptor by stromal progenitor cells during kidney development, as well as juxtaglomerular (JG) cells and glomerular mesangial cells in the adult kidney (Mochida et al., 2022, Hill et al., 2023). While our antibody labeling did not detect these PIEZO2^+^ cells, this was also noted in the study by Mochida et al. where RNAScope *in situ* hybridization was necessary to detect *Piezo2* expression. We have established the presence of PIEZO2^+^ sensory nerves in both the mouse and human kidney where they associate with the vasculature, glomeruli, and proximal tubules. We hypothesize that these sensory axons are poised to sense changes in pressure or volume within the kidney. It has been established that PIEZO2^+^ cells in the bladder communicate with pressure sensitive PIEZO2^+^ afferent nerves to mediate micturition (Marshall et al., 2020), and PIEZO2 signaling is critical for sexual function (Lam et al., 2023). We recently reported that PIEZO2 in JG cells modulates renin levels (Hill et al., 2023). However, further investigation is required to understand whether such communication between PIEZO2^+^ kidney cells and PIEZO2^+^ sensory neurons play a role in development or physiological function. Interestingly, many of these sensory nerves are seen looping in a “hairpin” fashion around glomeruli and tubules. Such anatomic position may better enable the ability to sense changes in pressure, volume, filtration rate, and tubule physiology ensuring homeostatic control. The looping may also enable innervation of structures within the kidney without the need for a singular synapse to each kidney cell. These findings further emphasize the need for studies investigating the physiological role of sensory afferent nerves in the control of kidney development and function in health and disease.

In addition to the sensory nerves, we also showed that TH^+^ sympathetic efferent nerves innervate the developing kidney. TH^+^ axons were detected as early as E14.5 and as development progressed, TH^+^ nerves associated with the vasculature, tubules, and glomeruli. Association with these structures correlates with the known physiological functions of sympathetic nerves in modulating vascular tone, tubular sodium reabsorption, and renin release by JG cells (Osborn et al., 2021; Osborn & Foss, 2017). It has been established that renal sympathetic nerves contribute to the pathogenesis of hypertension. Thus, procedures like catheter-based renal denervation are seen as novel therapies that could reduce blood pressure in hypertensive patients (Esler et al., 2012; Krum et al., 2013, 2014). However, this procedure non-selectively targets all renal nerves and with our observation of the significant overlap of sensory and sympathetic fibers innervating glomeruli, a deeper understanding of the interplay between sympathetic and sensory nerves in the mediation of physiology may inform efforts to optimize these ablation strategies.

Most visceral organs are innervated by both sympathetic and parasympathetic nerves. This is no exception to the kidney which was first hypothesized to harbor parasympathetic nerves that enter the kidney along the renal vasculature (Furness, 1999; Mitchell, 1950). Yet, there remains no significant anatomic evidence of extensive parasympathetic innervation within the kidney, and this remains a topic of debate. In our study, we observed TUBB3^+^ nerve fibers which colocalized with the cholinergic marker VAChT. However, this could represent a subset of immature sympathetic nerves expressing both VAChT and TH, although as development progressed, we observed VAChT^+^ nerves in the cortex associating with structures such as tubules which are more likely to represent mature, functional parasympathetic nerves. Higher resolution imaging studies are needed to determine if these are sympathetic nerves expressing cholinergic markers or whether they represent separate classes. Recently, research by Cheng et al. on adult mouse renal arteries showed the presence of periarterial cholinergic nerves which were separate from sympathetic and sensory nerve fibers (Cheng et al., 2022). Together, these findings point to the potential for greater contribution of parasympathetic activity in the modulation or kidney function than has been previously appreciated, although further characterization is necessary.

### Common DRG projections to organs of the urogenital system

The regulation of kidney homeostasis is in part maintained by neuronal modulation. Ablation of renal nerve activity is widely seen as a way to treat diseases like hypertension. Accurate mapping and understanding the molecular identity of kidney nerves will help guide such interventions. However, studies have largely focused on kidney sympathetic nerves and only recently has the contribution of sensory neurons to physiological regulation been more thoroughly investigated (Banek et al., 2016; Ong et al., 2019; Osborn et al., 2021). Building on these previous studies, we found that kidney sensory innervation is substantial and molecularly diverse. Axons from sensory neurons form an extensive network in the developing kidney that continues to expand with maturation. Validation of sensory innervation was provided by retrograde tracing and identified the DRG origins of these nerves. Kidney sensory innervation traces to the lower thoracic region from DRG T10-T13, the lumbar region from DRG L1-L6, and the sacral region from DRG S1 and S2. We cannot absolutely rule out some leakage of tracer into the pelvis and ureter, thus expanding the labeling to lower lumbar and sacral DRG, albeit at weaker levels. Despite this variable DRG labeling, the majority of axons innervating the kidney project from T10-13 and L1-L2. PIEZO2^+^ neurons similarly traced to the DRG at these levels. Tracing performed in the mouse bladder and prostate via the application of FastBlue dye revealed similar projection of neurons from T13, L1, L2, L6, and S1 (Lee et al., 2016). This suggests that organs of the genitourinary system are largely innervated from common DRG. Interestingly, our CTB tracing experiment also showed that contralateral DRG at L1 and L2 were positive for CTB. These data suggest that some axons innervating the kidney might originate from contralateral DRG or that the same DRG in the ipsilateral side sends axons to both kidneys. Concurrent retrograde labeling of sensory nerves in the bladder and colon showed that dichotomizing afferent nerves coming from an individual DRG neuron innervate both the colon and the bladder (Christianson et al., 2007). Such connectivity is important for integrating sensory input from multiple organs and likely plays a major role in conditions such as pelvic pain disorders.

### Establishing crosstalk during kidney development

During development the nerves lie in close association with the vasculature, tubules, and glomeruli suggesting potential functional crosstalk. Sympathetic and sensory nerves branch with the vasculature and dissociate to navigate through the tissue where they appear to loop around glomeruli and lie adjacent to tubules. Such close association likely allows for the formation of neuroeffector junctions with target structures and nerves to functionally innervate multiple structures as axons navigate around them. As reviewed previously (Barajas et al., 1984; DiBona & Kopp, 1997; Osborn et al., 2021), these fiber varicosities could impact adjacent kidney cells by establishing neuroeffector junctions more than 50μm away from the neurotransmitter release site, allowing for diffusion throughout the interstitium and to target cells. To investigate the presence of neuroeffector junctions in the kidney, we assessed the localization of synapsin I, which is a major component of pre-synaptic vesicles and important for axonogenesis (Chin et al., 1995). Early during development, we observed that synapsin I is mainly localized near vasculature and this localization profile is extended to the glomeruli and tubules by E18.5. Such findings agree with our current understanding of kidney development and physiology. Mouse kidney development continues postnatally contrary to human kidney development which is finished before birth (McMahon, 2016). We also know that the mouse kidney forms around 90% of its nephrons towards the end of ureteric branching with 40% of them differentiating after birth (Smyth, 2021). Yet, kidney function must be established very early on in newborn pups. Thus, it is likely important to establish proper innervation of the developing kidney to regulate homeostasis. Interestingly, data mining through publicly available single cell RNA-seq datasets of E18.5 kidneys (Combes et al., 2019) suggests that many AMPA (*Gria1*, *Gria3*, *Gria4*) and NMDA (*Grin3a*) receptors are expressed by different nephron and stromal progenitors, as well as early proximal tubules. While it has been established that glutamatergic signaling via NMDA maintains the epithelial phenotype of proximal tubular cells through calcium signaling in a proximal tubule cell line (HK-2) (Bozic et al., 2011), it remains unclear what role glutamate transport may play in the proper development of proximal tubules. Overall, the expression of neurotransmitter receptors and postsynaptic markers in the developing kidney suggests the potential for several neuronal signaling mechanisms that regulate development and function.

### Limitations of our studies

To our knowledge, our study is the first to provide a thorough characterization of kidney innervation detailing the spatial and temporal establishment of nerve networks during development, including association and putative crosstalk with target structures. Importantly, we establish that a significant portion of the nerve network consists of sensory afferent axons with the novel identification of PIEZO2^+^ mechanosensory neurons. However, it is important to note some assumptions which guided our study. For the classification of noradrenergic sympathetic nerves, we use TH as a marker. The strong expression of TH by sympathetic ganglion near the descending aorta support our finding that TH^+^ sympathetic nerves are abundant in the developing kidney. However, some TH^+^ axons in the developing kidney may project from the DRG (Meerschaert et al., 2020) thus we cannot omit that a certain percentage of these nerves may be sensory, although this is likely to be a minor percentage given the low level of TH expression within sensory ganglia prior to birth (Smith-Anttila et al., 2021). Additionally, the colocalization of TH and NPY in adults suggests that any significant sensory nerve production of TH is resolved by this point. Finally, we relied on the careful study of technical and biological replicates but did not quantify axon metrics such as density or percent of sympathetic versus sensory nerves aside from our adult glomerular quantitation. Hence, there is a possibility that we did not identify more subtle differences among stages and classes of nerves, in addition to sex differences. Future investigations are necessary to parse out these details and to also determine the significance of these nerves to overall kidney development and the establishment of proper function.

## Materials and methods

### Animals

All procedures and experiments were performed according to the guidelines of the National Institutes of Health Guide for the Care and Use of Laboratory Animals and were approved by the Institutional Animal Care and Use Committees of The University of North Carolina at Chapel Hill (protocols 19-183 and 22-136) and performed under the policies and recommendations of the International Association for the Study of Pain and approved by the Scripps Research Animal Care and Use Committee (protocol 08-0136). Mice were kept in standard housing with a 12-h light–dark cycle, with the room temperature kept around 22 °C, and humidity between 30% and 80% (not controlled). Mice were kept on pelleted paper (Scripps) or corncob (UNC) bedding and provided with paper square nestlets and polyvinyl chloride pipe (Scripps) or hut (UNC) enrichment with ad libitum access to food and water. Adult Swiss Webster (Taconic stock: SW MPF) and C57Bl6/J (Jackson Labs (JAX) stock #: 000664) mice were purchased from the respective vendor for experiments and in house matings. Swiss Webster incrosses were utilized for all experiments involving only embryonic and postnatal wild type mice. C57Bl/6J mice were utilized for maintenance of transgenic and knock-in lines and adult analyses. Mice utilized for adult analyses were 8 weeks to 4 months of age. *Trpv1^Cre^* (JAX stock #:017769)(Cavanaugh et al., 2011) were maintained as a homozygous line, *Pod-Cre^tg^* mice (JAX stock #:008205)(Moeller et al., 2003) were maintained as heterozygotes, *R26^tdTomato^* (JAX stock #:007914)(Madisen et al., 2010) mice were maintained as a homozygous line, and *Piezo2^EGFP-IRES-Cre^* (JAX stock #:027719) (Woo et al., 2014) mice were maintained as heterozygotes.

### Tissue collection and processing

For all mouse experiments, matings were set up in the afternoon and copulation plugs found the following day were designated as embryonic day (E)0.5. The day of birth was considered postnatal day (P)0. This occurred at E19.5, births earlier or later than E19.5 were not utilized in these studies. At the desired developmental time point, pregnant dams were sacrificed by CO_2_ inhalation, and death was confirmed by cervical dislocation. Mouse embryos were excised from the uterus and kidneys or whole urogenital systems dissected in ice-cold 1x phosphate-buffered saline (PBS). For genetic crosses, tail or ear clips were taken and incubated with DirectPCR lysis buffer (Viagen Biotech, 102-T) containing 10µg/mL proteinase K (Sigma, 3115836001), incubated at 55°C overnight, and denatured at 95°C for 20 min the next day. Genotyping was performed using JAX genotyping protocols and recommended primers with the following PCR parameters: annealing temperature T_anl_ = 58°C, elongation time t_elg_ = 40 sec for 35 cycles. To ensure biological significance, we analyzed a minimum of 3 animals per neuronal marker for both sexes at the following stages: E12.5, E13.5, E14.5, E16.5, E18.5, P0, P7, and adults (6-10 weeks old). For tissue processing, all steps below were performed on rotating shakers. Harvested kidneys or urogenital systems from embryonic and postnatal mice were fixed by immersion in fresh ice-cold 4% PFA-PBS (Fishersci, 15714) at 4°C for 30 min, while kidneys obtained from juvenile and adult mice were fixed by immersion in fresh ice-cold 4% PFA-PBS at 4°C for 2 hours. Next, all kidneys were washed in ice-cold 1x PBS three times for 10 min each at room temperature and stored at 4°C in 1x PBS until use. All kidneys destined to be sectioned were incubated in 30% sucrose-PBS solution at 4°C overnight. The next day, kidneys were then embedded in Tissue freezing media (Fishersci, 23730571) and sectioned at 12μm on a Leica cryostat CM1850. Slides were stored at -80°C until use.

Human kidney tissue samples were obtained from the UNC Tissue Procurement Facility and were de-identified and therefore were exempt from IRB approval. Tissue was obtained as a frozen block. A small piece of tissue was removed with a razor blade, fixed for 10 min in 4% paraformaldehyde, and subsequent processing for cryosectioning and immunofluorescence was carried out as for the mouse.

### Cholera Toxin B(CTB) and pAAV-FLEX-tdTomato retrograde tracing

Surgeries were performed as in Woodard et al., 2018. Briefly, mice were anesthetized with isoflurane inhalation and surgery started after all-paw reflexes disappeared. The right kidney was accessed via flank incision after the animal was shaved from shoulder to rump and flank to spine with a shaver designed for pet grooming. After appropriate cleaning of the surgical site, the skin was pinched with tweezers and a cut of approximately 1cm from the spine and below the ribcage was made with scissors. A similar cut was made on the muscle layer underneath the skin, the kidney was exposed and carefully isolated with cotton-tip applicators from peripheral fat or connective tissues in order to not disrupt nerve connections. Injections were made with a 10ul Hamilton syringe attached to a 32G needle. Tracers were injected at an approximate rate of 1µl/30sec and the needle was left inside the tissue after injection for an additional 1 minute to avoid back-leak. For retrograde labeling of the DRG innervating the kidney, 4 injections of 3µl 0.2% CTB-555 (Invitrogen, C34776) were administered to 4 males and 4 females on a C57Bl6/J and Swiss Webster background. For retrograde labeling of the PIEZO2^+^ nerves innervating the kidney, 4 injections of 5µl of AAV PHP.S (Addgene, 28306-PHP.S, *pAAV-FLEX-tdTomato*; deposited by Viviana Gradinaru, Chan et al. 2017) were administered to 2 males and 2 females which were *Piezo2^EGFP-IRES-Cre/+^*. All injections were administered directly into kidney tissue through the renal capsule. The 4 injections consisted of one cortical injection per pole of the kidney, one injection in the medulla, and one injection in the renal pelvis. After injections, the muscle and skin were closed with sutures, and bupivacaine was administered for analgesia. After suturing, mice were housed in a clean cage with heat pads underneath until ambulatory. Mice were euthanized 4 days after CTB injections and 4 weeks after AAV PHP.S injections. Mice were perfused with 30ml 1x PBS and 20ml 4% PFA-PBS with kidneys and DRG collected for tissue sectioning and imaging.

### Section immunofluorescence

Kidney sections for immunofluorescence were removed from the -80°C freezer and allowed to equilibrate to room temperature for 10 min. Sections were rehydrated, and the freezing medium was removed by incubating the slide in 1x PBS for 10 min in a Coplin jar. After rehydration, excess PBS was removed from the slide then a hydrophobic barrier was drawn around the section using a hydrophobic pap pen (Vector Laboratories, Cat#H-4000). Sections were incubated for 30 min in blocking solution (ice-cold PBS solution with 4% donkey serum (Equitech-Bio, SD30-0100), 1% bovine serum albumin (Fishersci, BP9706-100), and 0.25% Triton X-100 (Fishersci, BP151-500). Slides were then incubated at room temperature for 2 hours covered from light with primary antibodies in block solution (see Supplemental Table 1 for primary antibodies utilized; PIEZO2 antibodies were combined as a mixture for optimal detection although each work independently as validated by co-immunostaining with GFP on DRG isolated from *Piezo2^EGFP-IRES-Cre/+^* mice ). After primary incubation, slides were washed 3 times for 5 min at room temperature covered from light with wash solution (PBS + 0.25% Triton X-100) before incubating them for 45 min at room temperature covered from light with the appropriate Alexa Fluor secondary antibody (Life Technologies, 488, 568, or 647 conjugated; 1:1000) in blocking solution. Finally, slides were washed 4 times for 5min at room temperature covered from light using wash solution with the last wash containing DAPI diluted at 1:10000. Slides were mounted using Prolong gold mounting media (ThermoFisher, P36930), and images were captured with a Zeiss 880 confocal laser scanning microscope.

For the NFH, NPY/TH and CGRP/TH immunofluorescence, C57BL6/J adult mice were euthanized via isoflurane overdose and transcardially perfused with 20 mL ice-cold PBS pH 7.4 followed by 30 mL ice-cold 4% PFA in PBS. Kidneys were dissected and fixed overnight in 4% PFA at 4°C. Kidneys were then rinsed in PBS and cryoprotected in 30% sucrose for 24-48 hours at 4°C prior to embedding and freezing in OCT medium (Sakura). 20 µm sections were prepared on glass slides using a cryostat. Tissues were allowed to air-dry for 30 minutes at room temperature and hydrophobic barriers were drawn using a PAP pen (Vector Biolabs, H-4000). Slides were washed in PBS and blocked in 5% Normal Donkey Serum (for NPY IHC, Sigma-Aldrich, D9663) or Normal Goat Serum (all other experiments, Life Technologies, PCN5000) in PBS with 0.3% Triton X-100 (PBST) for two hours at room temperature in a humidified chamber. Slides were incubated overnight at 4°C in the following primary antibodies in 0.1% PBST: CGRP, TH (chicken), NFH, or NPY. Slides were washed in 0.3% PBST followed by PBS and then incubated for two hours at room temperature in the following secondary antibodies in 0.1% PBST: donkey anti-goat 647 (Life Technologies, A21447, 1:1000), donkey anti-chicken 488 (Life Technologies, A78948, 1:1000), goat anti-Rabbit 647 (Life Technologies, A21245, 1:1000), goat anti-chicken 488 (Life Technologies, A11039, 1:1000). Donkey secondary antibodies were used for NPY staining experiments and goat secondary antibodies were used for all other experiments. Samples were mounted using Fluoromount G + DAPI (Life Technologies, 00-4959-52) and sealed with No. 1.5 coverslips and clear nail polish and stored at 4°C. Samples were imaged on a Nikon AX confocal microscope.

### CGRP/TH glomeruli quantification

Maximum intensity z-projections were generated and glomeruli were identified from the DAPI signal as regions of interest (ROI). Each individual ROI was manually assessed for the presence of TH or CGRP immunoreactivity at the Bowman’s capsule or at the juxtaglomerular region adjoining the glomerulus and the arterioles. ROI with immunoreactivity at either location were determined to be innervated. For the TH staining, substantial non-neuronal tubular immunoreactivity was observed that was readily distinguishable from nerve fibers. The tubular immunoreactivity was not used to quantify innervation. FIJI (2.0.0) and Prism 10 (10.1.1) software were utilized for analyses.

### Wholemount immunofluorescence and tissue clearing

For mouse embryonic and postnatal kidneys or urogenital systems (E12.5-P7), we used a modified version of the original iDISCO protocol (Renier et al., 2014) for wholemount immunolabelling (see Supplemental Table 2) using validated antibodies and concentrations indicated in Supplemental Table 1. All steps are performed on rotating shakers. We first incubated whole pre-fixed tissue in 1ml blocking solution (ice-cold PBS solution with 10% (v/v) Heat Inactivated Sheep Serum (HISS) (ThermoFisher, 16070096) and 0.5% (v/v) Triton X-100) for 1hr at room temperature. Then, we incubated the blocked kidney in 1ml primary antibody in block solution for 7 days at 4°C while covered from light. The labeled kidney was then washed using 1ml wash buffer (PBS solution with 0.25% Triton X-100) for a whole day at room temperature covered from light with wash solution changed every 1 hour. After washing, labeled kidneys were incubated in 1ml secondary antibody in block solution at a concentration of 1:1000 for 3.5 days at 4°C while covered from light. After secondary antibody labeling, kidneys were washed again using 1ml wash buffer for a whole day at room temperature covered from light with wash solution changed every 1 hour. Before optical clearing, successful labeling was checked on a widefield Leica fluorescent microscope (Leica DMi8). Labeled kidneys were then embedded in 1% agarose (Fishersci, BP160-500) gel made with 1x TAE, the agarose block was cut to a size of 5mmx5mm to ensure proper mounting on the light-sheet microscope. To clear kidneys, we first dehydrated the agarose block in a methanol-H_2_0 (Fishersci, A433P-4) series wash at room temperature (25%, 50%, 75%, and 100%) for 1 hour each minimum covered from light. Next, dehydrated agarose blocks were incubated in 66% v/v dichloromethane (DCM) (Sigma, 270997-2L)/33% v/v methanol for 3 hours at room temperature covered from light. Following this, the agarose blocks were then incubated in 100% DCM for 15min x 2 at room temperature covered from light. Finally, the fully dehydrated agarose blocks were incubated in 100% v/v dibenzyl ether (DBE) (Sigma, 33630-1L) and stored at room temperature away from light. The blocks became transparent and invisible within 5-7hr and were stored in DBE until imaging. The DBE was changed out for fresh DBE before imaging the sample.

### 3D light-sheet and confocal imaging

All whole-mount images were obtained using the LaVision BioTec Ultramicroscope II light-sheet system at the UNC Microscopy Services Laboratory (MSL). This light-sheet is outfitted with an Andor Zyla 5.5 sCMOS detector, an Olympus MVPLAPO 2X/0.5 objective coupled to a zoom body ranging from 1.26x to 12.6x with a corrected dipping cap and a working distance of 5.7mm. All whole-mount images in this study were imaged with laser wavelengths: 488, 561, and 647nm as well as 3 angled light-sheet from one side only with the following parameters: sheet NA of 0.016, horizontal focus set in the middle of the sample, correction collar set at 3.5 and 2µm Z-spacing (step size). All tiff images are generated in 16-bit. All confocal images were obtained using a Zeiss 880 confocal laser scanning microscope in the UNC Hooker Imaging Core via the Zen 2 software with 34-channel gallium arsenide phosphide (GaAsP) detectors. All images obtained were imaged using a Plan-Neofluar 40x/NA:1.3 or 63x/NA:1.4 oil objective with the following working distance respectively: 0.21mm and 0.19mm. Pinhole size of 1AU set on the longest wavelength fluorophore with the same diameter size for all the rest. All image acquisition settings can be found in Supplemental Table 3.

### Image processing

Images, 3D volume, and movies were generated using Imaris x64 software version 9.9.1. All stack images obtained from light-sheet imaging were first converted into an imaris file (.ims) using Imaris File Converter version 9.8.2. All wholemount images and videos were obtained using the “snapshot” and “animation” tools in Imaris 9.9.1. files obtained from Zeiss confocal microscope (.czi) were converted into an Imaris file using Imaris File Converter 9.8.2. Tilescan images were stitched with 20% overlapped using Imaris stitcher 9.9.1 and image analysis was done in Imaris version 9.9.1.

### Receptor and postsynaptic marker scRNA-seq enrichment

To identify the kidney cell populations in which nerve associated receptors and markers were expressed, the selected genes were analyzed for their relative enrichment within the single cell RNA-seq data from E18.5 kidneys (Combes et al., 2019). Heatmaps were generated in R (R Core Team, 2021) using the ggplot2 package (Wickham, 2009) and ComplexHeatmap (Gu et al. 2016). Log fold change values were obtained from Combes et al., 2019 Table S1 “lookup” tab.

## Acknowledgments

We thank members of the O’Brien Lab for their critical insights throughout the project. We would like to thank Dr. Pablo Ariel director of the UNC Microscopy Services Laboratory for expert training in light-sheet image acquisition and analysis and lending his expertise throughout the project, and Dr. Wendy Salmon director of the UNC Hooker Imaging Core for confocal microscopy imaging training. The Microscopy Services Laboratory in the Department of Pathology and Laboratory Medicine and the Department of Cell Biology and Physiology Hooker Imaging Core are supported by the NIH P30CA016086 Cancer Center Core Support Grant to the UNC Lineberger Comprehensive Cancer Center. Light-sheet microscopy at MSL is also supported in part by the North Carolina Biotech Center Institutional Support Grant 2016-IDG-1016, and Andor Dragonfly microscope was funded with support from National Institutes of Health grant S10OD030223.

## Authors Contributions

L.L.O. conceptualized the study, supervised research, designed and guided experiments, interpreted data, helped assemble figures, and revised the manuscript. P-E.Y.N. helped design and performed experiments, data analysis, interpreted data, assembled figures, and wrote the manuscript. S.R.M. performed mouse surgery required for retrograde tracing, helped guide experiments, and contributed to the Piezo2 analysis. A.T. helped inject mice for retrograde tracing experiment. A.T. and M.G. dissected, imaged and analyzed DRGs data. G.S. interpreted retrograde tracing data and supervised A.T. and M.G. research. R.Z.H. performed NFH, NPY/TH, and CGRP/TH immunostainings, interpreted the results, and assembled the associated figures. S.S. immunostained and quantified the CGRP/TH innervation of glomeruli. R.Z.H. supervised S.S. All authors contributed to the manuscript text relevant to their contributions.

---

**Figure S1:**
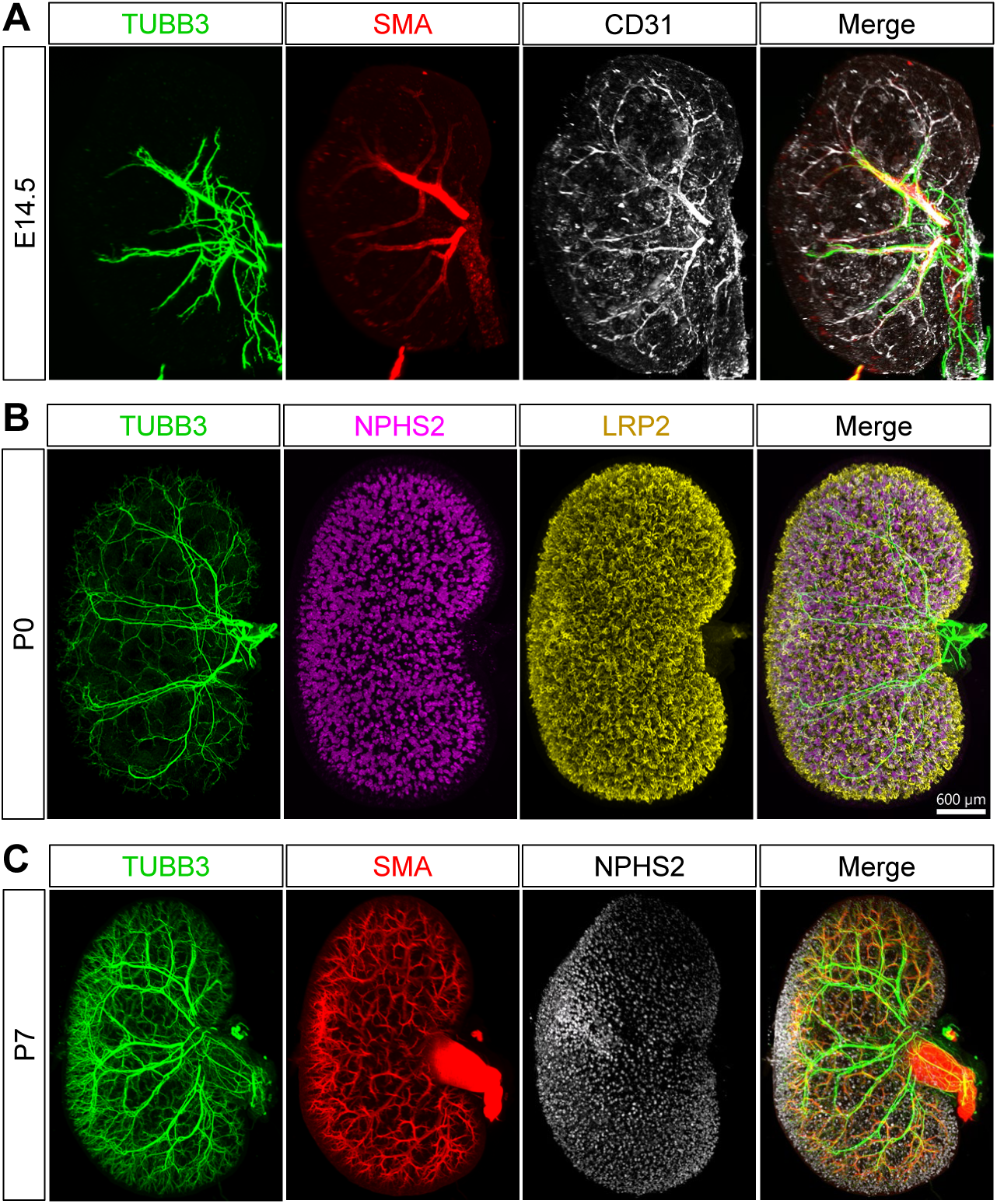
Innervation of the embryonic and postnatal kidney. **(A)** Wholemount immunostain of an E14.5 kidney for TUBB3 (green), SMA (red), and CD31 (grey). Axons track closely with the developing arteries despite the presence of an extensive CD31+ endothelial network. (**B)** Wholemount immunostain of a P0 kidney for TUBB3 (green), NPHS2 (podocytes, magenta), and LRP2 (proximal tubules, yellow). The neuronal network continues to expand and navigate towards and around kidney structures. **(C)** Wholemount immunostain of a P0 kidney for TUBB3 (green), SMA (red), and NPHS2 (podocytes, grey). The neuronal network has continued to grow as the vasculature expands and the last wave of nephrons differentiate.

**Figure S2:**
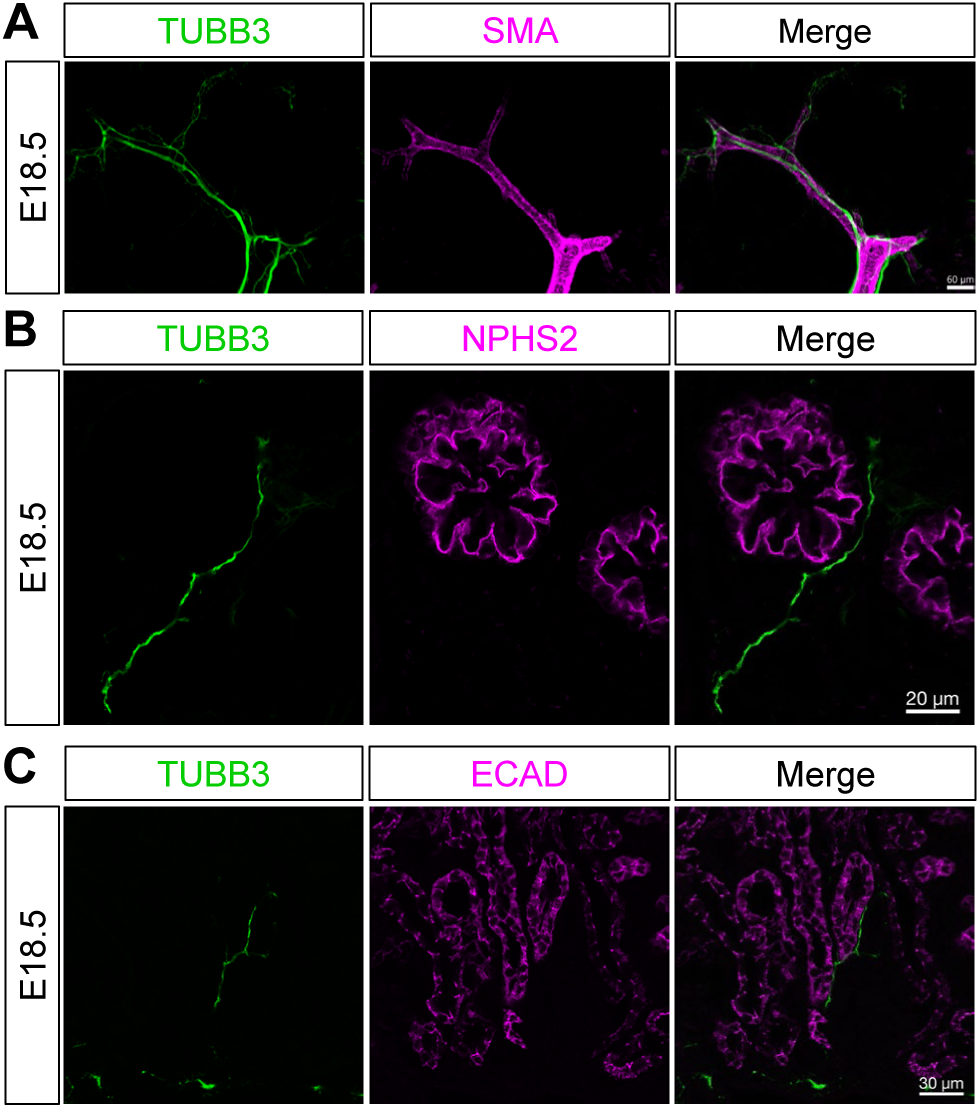
Axons associate with kidney structures during embryonic development. **(A)** Image of an E18.5 kidney section immunostained for vasculature (SMA, magenta) and nerves (TUBB3, green). Axons are tightly associated with the SMA+ vasculature. **(B)** Image of an E18.5 kidney section immunostained for nerves (TUBB3, green) and glomeruli (NPHS2, magenta). Axons approach the glomeruli and associate with the Bowman’s capsule as they navigate through the kidney. **(C)** Image of an E18.5 kidney section immunostained for nerves (TUBB3, green) and tubule epithelium (ECAD, magenta). Axons are observed closely associating with ECAD^+^ tubules. A minimum of n=3 kidneys were analyzed for each stage.

**Figure S3:**
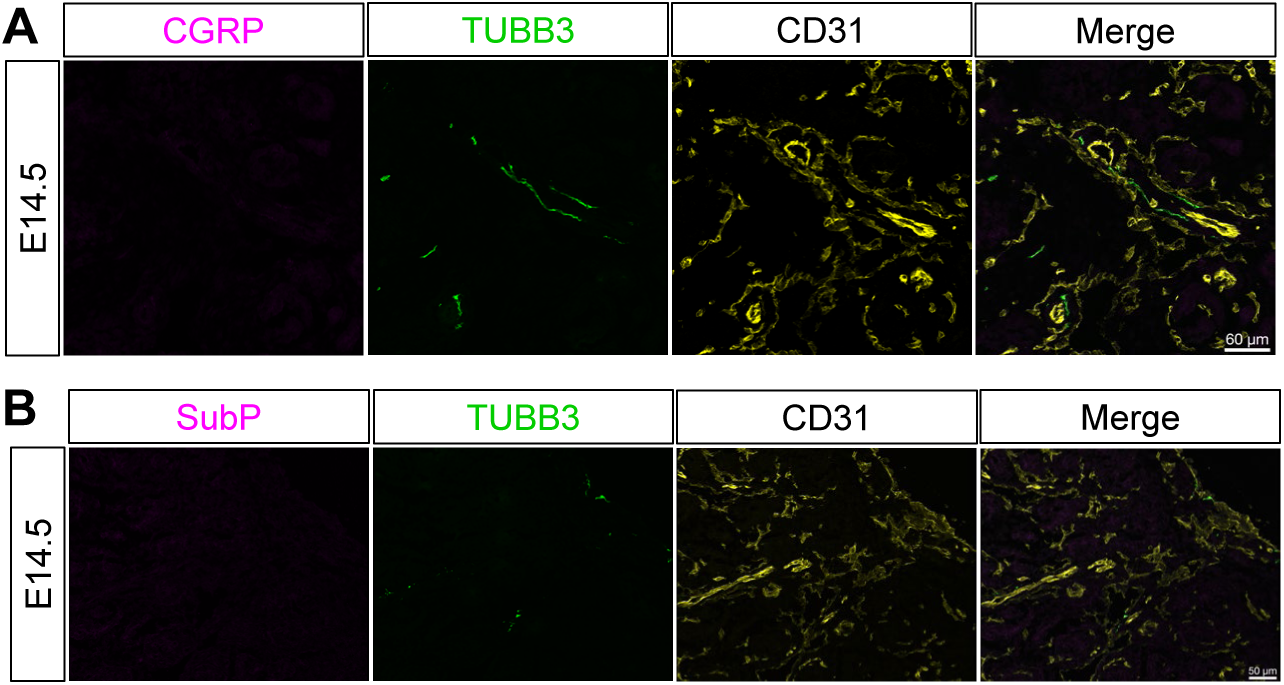
Kidney peptidergic sensory innervation is delayed. **(A)** E14.5 kidney sections immunostained for CGRP (magenta) shows that axons at this stage (TUBB3, green) that primarily associate with the vasculature (CD31, yellow) are not yet CGRP+. (B) E14.5 kidney sections immunostained for substance P (SubP, magenta) shows that axons at this stage (TUBB3, green) that primarily associate with the vasculature (CD31, yellow) are not yet SubP+.

**Figure S4:**
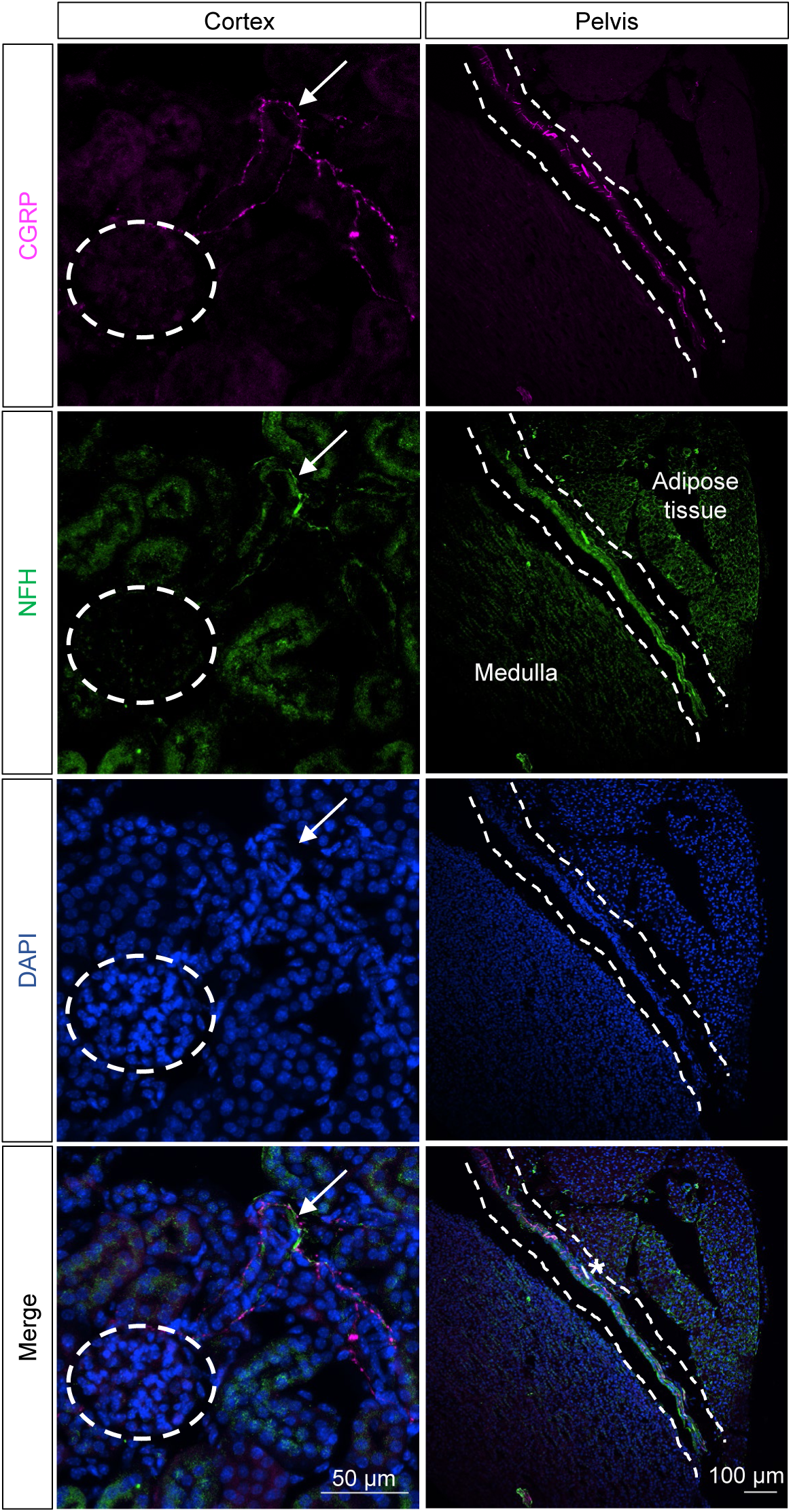
Myelinated NFH+ fibers sparsely innervate the kidney cortex and pelvis. Image of the adult kidney cortex (left panel) immunostained for the CGRP (magenta), the cytoskeletal protein Neurofilament heavy polypeptide (NFH, green), and the nuclear counterstain DAPI (blue). A glomerulus is encircled in a white dotted line and the arrow indicates a single NFH+ fiber that does not approach the glomerulus and which is negative for CGRP immunoreactivity. These cortical NFH+ fibers were rarely observed. Image is representative of n=2 male mice. Image of the adult renal pelvis (right panel) immunostained for the CGRP (magenta), NFH (green), and DAPI (blue). The border of the pelvic urothelium is outlined with a white dashed line. NFH+ fibers were observed within the urothelium and occasionally these appeared to overlap with CGRP+, however, the density of CGRP immunoreactivity in the pelvis made it challenging to distinguish individual fibers. The asterisk (*) denotes a portion of nerve trunk comprised of multiple fibers containing both CGRP and NFH immunoreactivity. Image is representative of n=2 male mice.

**Figure S5:**
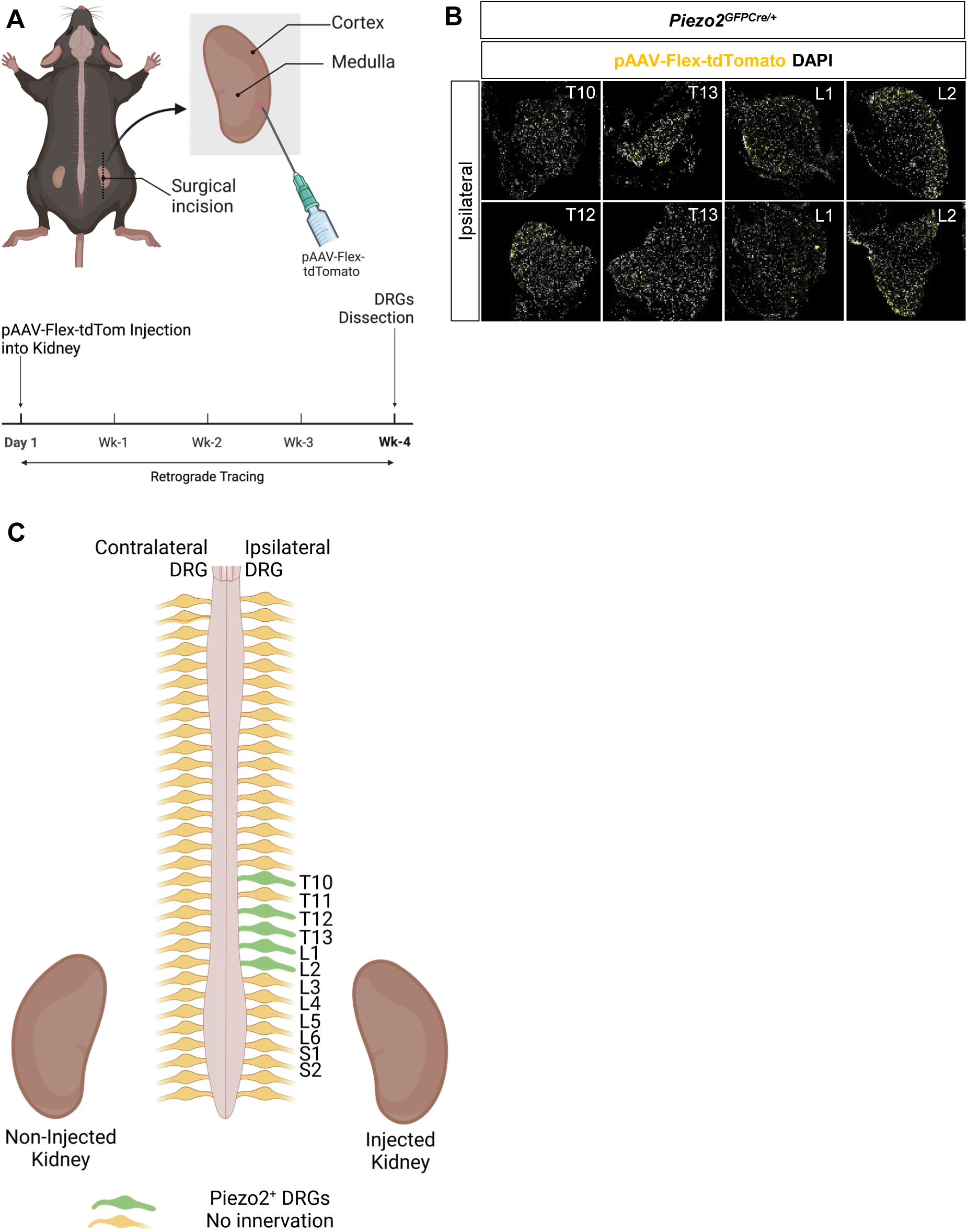
pAAV-Flex retrograde tracing of PIEZO2^+^ sensory nerves show labeling of lower thoracic and upper lumbar DRGs. **(A)** Schematic of retrograde tracing injections and timeline. pAAV-Flex-tdTomato was injected into the right kidney of *Piezo2^GFPCre/+^* mice. Three weeks following ipsilateral (injected side) and contralateral (non-injected side) DRGs were collected for analysis. **(B)** PIEZO2^+^ renal sensory innervation primarily traces in the ipsilateral DRG in the lower thoracic region (T10-T13) and the upper lumbar region (L1-L2). Scale = 50μm. **(C)** Schematic representation of the spinal level and DRG origin of kidney PIEZO2+ sensory afferent innervation.

**Figure S6:**
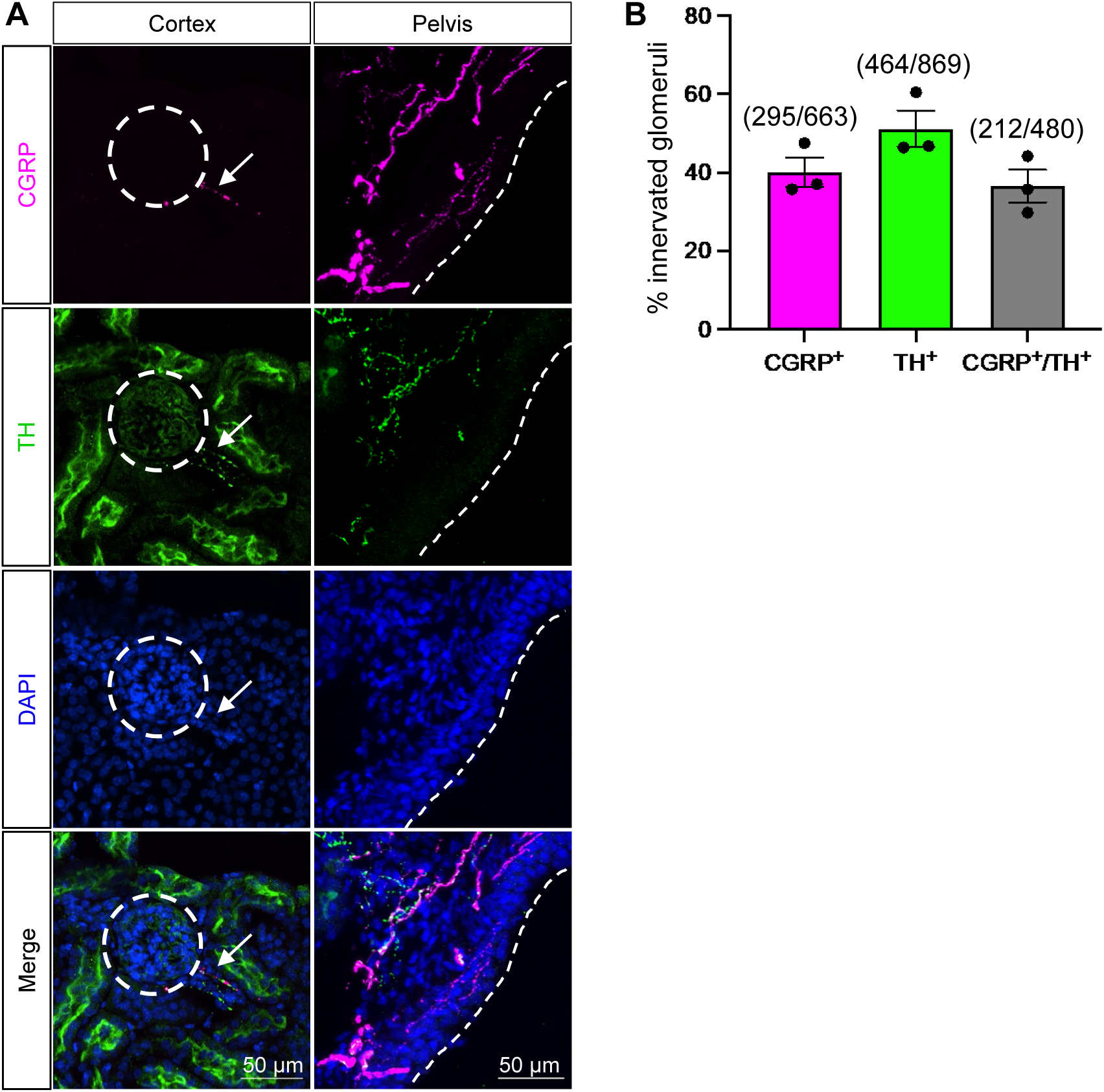
Analysis of sensory and sympathetic innervation of glomeruli and the renal pelvis. **(A)** Image of the adult kidney cortex (left) and the renal pelvis (right) immunostained for CGRP (magenta), TH (green), and the nuclear counterstain DAPI (blue). A glomerulus is encircled in a white dotted line and the border of the pelvic urothelium is outlined with a white dashed line. There is both sensory and sympathetic innervation of the arterioles adjacent to the glomerulus (white arrow) and the renal pelvis. Sensory nerves in the renal pelvis project more superficially into the urothelium than the sympathetic fibers. **(B)** Quantification of sensory and sympathetic innervation of glomeruli from n=3 animals (2 male and one female) expressed as a percentage of the total glomeruli. Total number of glomeruli analyzed is indicated above each bar. Error bars represent mean +/- SEM.

**Figure S7:**
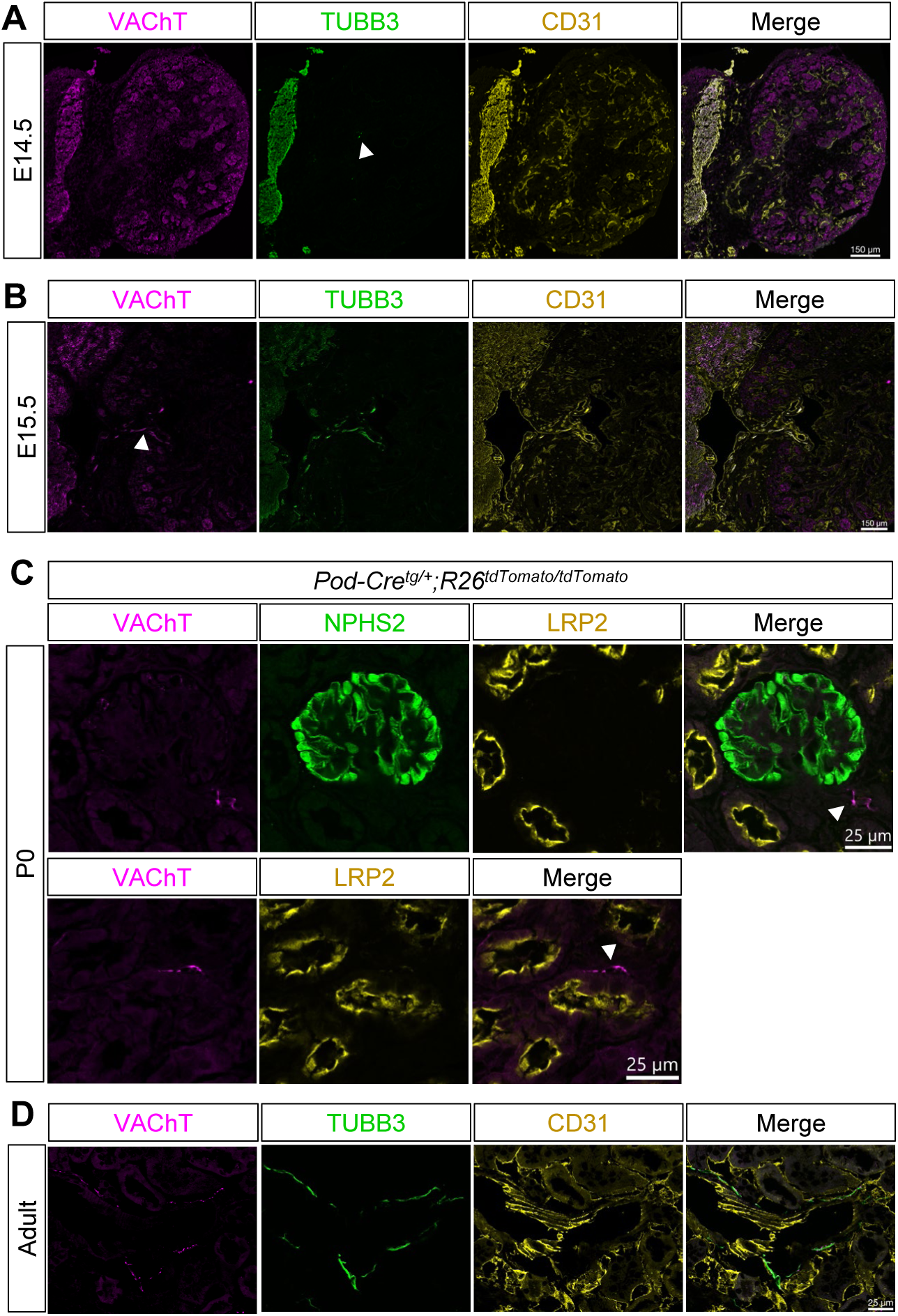
The cholinergic marker VAChT labels axons in the developing kidney. **(A)** E14.5 kidney sections immunostained for VAChT (magenta), TUBB3 (green), and CD31 (endothelium) show that axons (white arrowhead) are not positive for VAChT at this stage. VAChT signal is primarily background, non-specific signal. **(B)** E15.5 kidney sections immunostained for VAChT (magenta), TUBB3 (green), and CD31 (endothelium) show that VAChT signal (white arrowhead) is similar to TUBB3 axonal signal (green). **(C)** Sections of *Pod-Cre^tg/+^;R26^tdTomato/tdTomato^* P0 kidneys immunostained for VAChT (magenta) show VAChT+ axons near glomeruli (tdTomato, green; white arrowheads, top panel) and proximal tubules (LRP2, yellow; White arrowhead, bottom panel). **(D)** Adult kidney sections immunostained for VAChT (magenta), TUBB3 (green), and CD31 (endothelium) show that VAChT signal overlaps with TUBB3 and closely associates with the vasculature.

**Figure S8:**
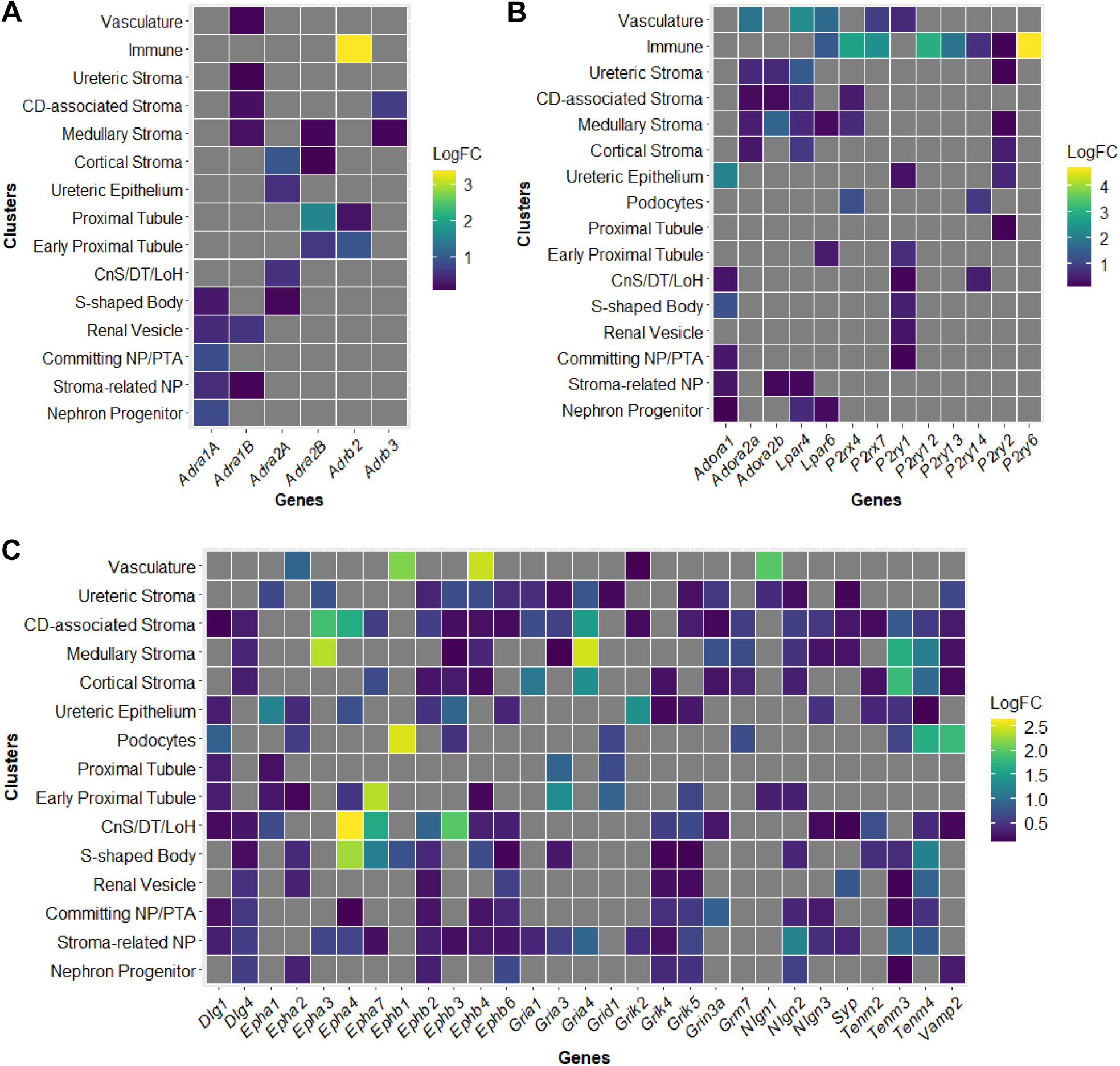
Differential enrichment of adrenergic, purinergic, and post-synaptic markers within the different cell clusters of the E18.5 kidney. **(A)** Heatmaps showing relative enrichment of adrenergic receptors in the different kidney cell types (clusters) from E18.5 whole kidney scRNA-seq data (Combes et al., 2019). **(B)** Heatmaps showing relative enrichment of purinergic receptors in the different kidney cell types (clusters) at E18.5. **(C)** Heatmaps showing relative enrichment of post-synaptic markers in the different kidney cell types (clusters) at E18.5. The scale indicates log fold change relative to the other cell clusters. NP=nephron progenitor, PTA = pretubular aggregate, DT = distal tubule, CD = collecting duct, LoH=Loop of Henle, CnS=connecting segment. LogFC = log fold change differential expression.

**Suppl. Table 1:** Wholemount tissue processing steps for iDISCO^+^ and light-sheet microscopy.

| Steps |  | Modality | Time | Temperature | Volume (mL) |
| --- | --- | --- | --- | --- | --- |
| 1 | Blocking:<br>PBS, 10% (v/v) HISS, 0.5% (v/v) Triton-x 100 | passive perfusion<br>on rocker cover from light | 1hr | RT | 1mL/Kidney pair |
| 2 | Primary Antibody Solution:<br>PBS, 10% (v/v) HISS, 0.5% (v/v) Triton-x 100, Primary antibody | passive perfusion<br>on rocker cover from light | 7 days | 4°C | 1mL/Kidney pair |
| 3 | Wash Solution:<br>PBS, 0.25% (v/v) Triton-x 100 | passive perfusion<br>on rocker cover from light | 1 days | RT | 1mL/Kidney pair |
| 4 | Secondary Antibody Solution:<br>PBS, 10% (v/v) HISS, 0.5% (v/v) Triton-x 100, Secondary antibody | passive perfusion<br>on rocker cover from light | 3.5 days | 4°C | 1mL/Kidney pair |
| 5 | Wash Solution:<br>PBS, 0.25% (v/v) Triton-x 100 | passive perfusion<br>on rocker cover from light | 1 days | RT | 1mL/Kidney pair |
| 6 | Embed kidneys in 1% Agarose gel<br>to a 5x5mm block | n/a |  |  |  |
| 7 | Methanol Dehydration:<br>25% (v/v) Methanol in water | passive perfusion<br>on rocker cover from light | 1hr | RT | 5mL/Kidney pair |
| 8 | Methanol Dehydration:<br>50% (v/v) Methanol in water | passive perfusion<br>on rocker cover from light | 1hr | RT | 5mL/Kidney pair |
| 9 | Methanol Dehydration:<br>75% (v/v) Methanol in water | passive perfusion<br>on rocker cover from light | 1hr | RT | 5mL/Kidney pair |
| 10 | Methanol Dehydration:<br>100% (v/v) Methanol | passive perfusion<br>on rocker cover from light | 1hr | RT | 5mL/Kidney pair |
| 11 | DCM Dehydration:<br>66% (v/v) DCM/33% (v/v) Methanol | passive perfusion<br>on rocker cover from light | 3hrs | RT | 5mL/Kidney pair |
| 12 | DCM Dehydration:<br>100% (v/v) DCM | passive perfusion<br>on rocker cover from light | 15min | RT | 5mL/Kidney pair |
| 13 | DCM Dehydration:<br>100% (v/v) DCM | passive perfusion<br>on rocker cover from light | 15min | RT | 5mL/Kidney pair |
| 14 | 100% (v/v) DBE Incubation | cover from light | Indefinitely | RT | 5mL/Kidney pair |

**Suppl. Table 2:**
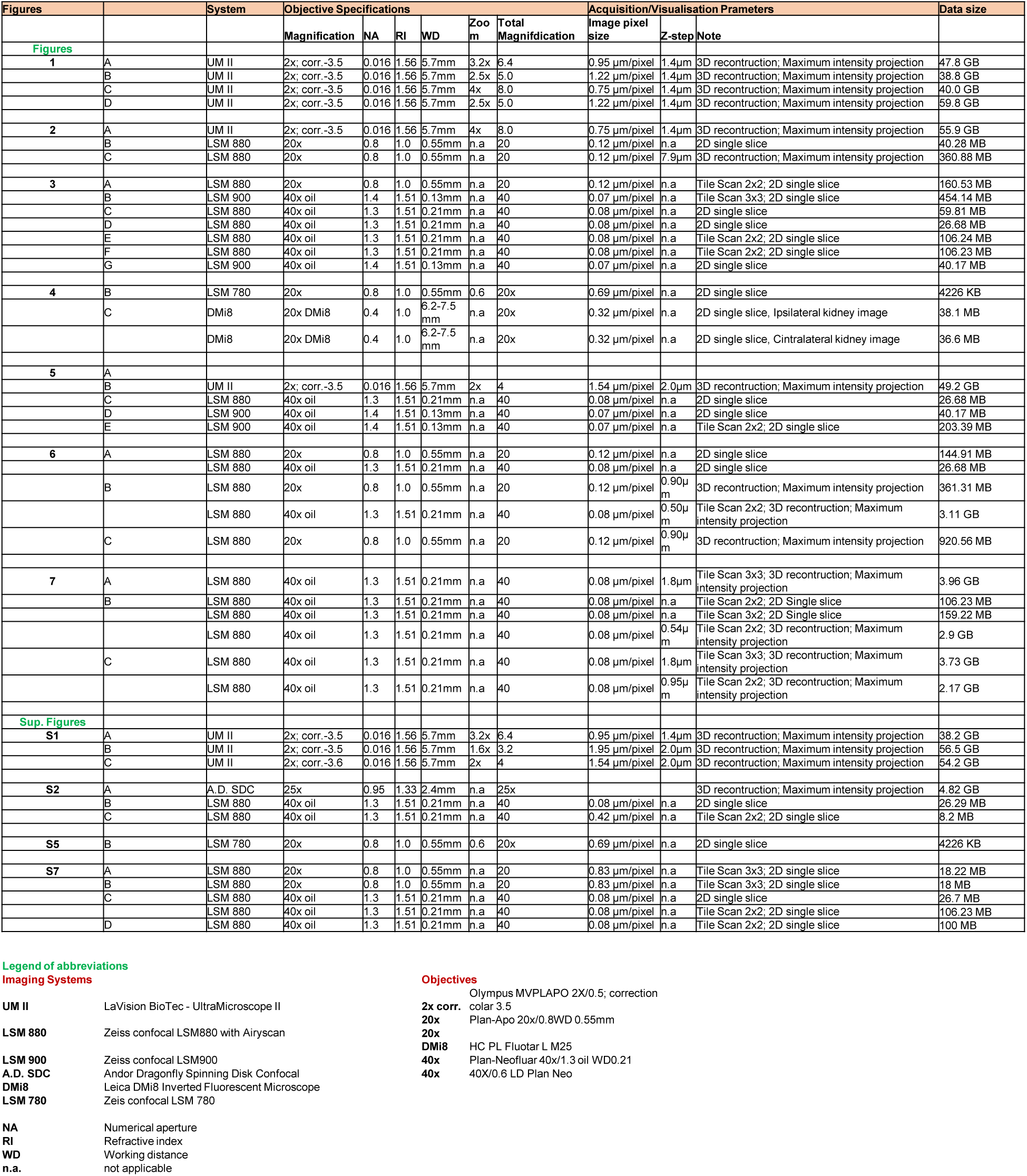
Imaging parameters utilized in this study.

**Suppl. Table 3:** List of primary antibodies used in this study.

| Antibody | Vendor | Cat # | Host | Dilution IF | Dilution WM | Alternate name, labels |
| --- | --- | --- | --- | --- | --- | --- |
| CD31 | BD Pharmingen | 550274 | Rat | 1:100 | 1:50 | Pecam1, Endothelial cell |
| CGRP | Immunostar | 24112 | Rabbit | 1:250 | n/a | Calcitonin Gene-Related Peptide, Peptidergic sensory nerve |
| CytoK | Sigma-Aldrich | C2931-.2ML | Mouse IgG1 | 1:500 | 1:250 | Cytokeratin, Ureteric tree |
| ECAD | Invitrogen | 131900 | Rat | 1:500 | 1:250 | E-cadherin, Tubule epithelium |
| LRP2 | MyBiosource | MBS690201 | Mouse IgG1 | 1:500 | 1:250 | Megalin, Proximal tubule |
| NFH | Abcam | AB4680 | Chicken | 1:1000 | n/a | Neurofilament heavy polypeptide, |
| NPHS2 | Invitrogen | PA579757 | Rabbit | 1:500 | 1:250 | Podocin, Podocytes, Glomeruli |
| NPY | Novus Biologicals | NBP1-46535SS | Chicken | 1:250 | n/a | Neuropeptide Y, Sympathetic nerve |
| PIEZO2 | Novus Biologicals | NBP1-78624 | Rabbit | 1:250 | n/a | Mechanosensitive sensory nerve |
| PIEZO2 | Novus Biologicals | NBP1-78538 | Rabbit | 1:250 | n/a | Mechanosensitive sensory nerve |
| PODXL | R&D Systems | AF1658 | Goat | 1:500 | 1:250 | Podocalyxin, glomeruli, endothelium |
| SMA-488 | Invitrogen | 53-9760-82 | Mouse IgG2a | 1:500 | 1:250 | Smooth muscle actin, Arterial tree |
| SubP | Immunostar | 20064 | Rabbit | 1:100 | n/a | Substance P<br>Peptidergic sensory nerve |
| SYN1 | Invitrogen | 51-5200 | Rabbit | 1:1000 | n/a | Synapsin I,<br>Synaptic vesicle marker |
| TH | Milipore Sigma | AB152 | Rabbit | 1:500 | 1:250 | Tyrosine Hydroxylase<br>Noradrenergic sympathetic nerve |
| TH | Abcam | AB76442 | Chicken | 1:250 | n/a | Tyrosine Hydroxylase<br>Noradrenergic sympathetic nerve |
| TUBB3 | BioLegend | 801202 | Mouse IgG2a | 1:500 | 1:250 | Class 3 Tubulin<br>Pan-neuronal marker |
| TUBB3-594 | BioLegend | 801208 | Mouse IgG2a | 1:500 | 1:250 | Class 3 Tubulin<br>Pan-neuronal Marker |
| VACHT | SySy | 139105 | Guinea-Pig | 1:100 | n/a | Vesicular acetylcholine transporter,<br>Parasympathetic (Cholinergic) nerve |
| RFP | Rockland | 600901379 | Chicken | 1:500 | 1:250 | Recombinant RFP (tdTomato) |

**Movie S1: Movie associated with Fig. 1A**

**Movie S2: Movie associated with Fig. 1B**

**Movie S3: Movie associated with Fig. 1B**

**Movie S4: Movie associated with Fig. 1C**

**Movie S5: Movie associated with Fig. 1D**

**Movie S6: Movie associated with Fig. 8A**

**Movie S7: Movie associated with Fig. 8C**

